# When inefficient speech-motor control affects speech comprehension: atypical electrophysiological correlates of language prediction in stuttering

**DOI:** 10.1101/2021.10.28.466231

**Authors:** Simone Gastaldon, Pierpaolo Busan, Giorgio Arcara, Francesca Peressotti

**Author notes:** **Corresponding authors:** Simone Gastaldon, Francesca Peressotti, Address: Dipartimento di Psicologia dello Sviluppo e della Socializzazione (DPSS), University of Padova, Via Venezia 8, 35131, Padova (PD), Italy. **Competing Interest Statement:** No competing interests to declare. **DISCLAIMER:** This is a provisional pre-peer-review manuscript. We invite interested readers to check the manuscript page on *bioRxiv* for any updated versions and for future notification of publication in a peer-reviewed journal.

## Abstract

It is well attested that people predict forthcoming information during language comprehension. The literature presents different proposals on how this ability could be implemented. Here, we tested the hypothesis according to which language production mechanisms have a role in such predictive processing. To this aim, we studied two electroencephalographic correlates of predictability during speech comprehension ‒ pretarget alpha‒beta (8-30 Hz) power decrease and the post-target N400 event-related potential (ERP) effect, ‒ in a population with impaired speech-motor control, i.e., adults who stutter (AWS), compared to typically fluent adults (TFA). Participants listened to sentences that could either constrain towards a target word or not, allowing or not to make predictions. We analyzed time-frequency modulations in a silent interval preceding the target and ERPs at the presentation of the target. Results showed that, compared to TFA, AWS display: i) a widespread and bilateral reduced power decrease in posterior temporal and parietal regions, and a power increase in anterior regions, especially in the left hemisphere (high *vs*. low constraining) and ii) a reduced N400 effect (non-predictable *vs*. predictable). The results suggest a reduced efficiency in generating predictions in AWS with respect to TFA. Additionally, the magnitude of the N400 effect in AWS is correlated with alpha power change in the right pre-motor and supplementary motor cortex, a key node in the dysfunctional network in stuttering. Overall, the results support the idea that processes and neural structures prominently devoted to speech planning and execution support prediction during language comprehension.

**Significance Statement:** The study contributes to the developing enterprise of investigating language production and comprehension not as separate systems, but as sets of processes which may be partly shared. We showed that a population with impaired speech-motor control, i.e., adults who stutter, are characterized by atypical electrophysiological patterns associated with prediction in speech comprehension. The results highlight that an underlying atypical function of neural structures supporting speech production also affects processes deployed during auditory comprehension. The implications are twofold: on the theoretical side, the study supports the need for a more integrated view of language comprehension and production as human capabilities, while on the applied and clinical side, these results might open new venues for efficient treatments of developmental stuttering.

## Introduction

Traditionally, research on human language has tackled production and comprehension as separate systems, occasionally interacting with each other but fundamentally distinct (Pickering & Garrod, 2013). This separation was mainly supported by research in classic aphasiology, showing asymmetries in understanding, producing, and repeating words and sentences in brain-lesioned patients (Shalom & Poeppel, 2008), and research on language acquisition, showing asymmetries in the developmental trajectories of comprehension and production skills in children (Clark & Hecht, 1983; Hendriks & Koster, 2010). However, these asymmetries may not necessarily reflect the separation between two distinct systems (Keenan, 1968; Tremblay & Dick, 2016; Chater, McCauley, & Christiansen, 2016). Furthermore, a growing body of research shows that the neural substrates underlying comprehension and production largely overlap (AbdulSabur et al., 2014; Giglio et al., 2021; Lukic et al., 2020; Silbert et al., 2014; Walenski et al., 2019), supporting the view that comprehension and production draw from processes and representations that are at least partially shared (Chater et al., 2016; Dell & Chang, 2014; Fairs et al., 2021; Gambi & Pickering, 2017; McQueen & Meyer, 2019; Pickering & Garrod, 2013).

In this study we focused on one aspect that is central to many integrated approaches, i.e., prediction. Language comprehension is the result of both bottom-up and top-down processes. Comprehenders not only incrementally integrate incoming information but also actively predict (i.e., pre-activate) different features of what they are likely to encounter by exploiting multiple cues and pieces of information (Kuperberg & Jaeger, 2016). Many researchers – albeit with partially different approaches – proposed that prediction processes during comprehension are linked to processes and representations traditionally attributed to language production (Christiansen & Chater, 2016; Dell & Chang, 2014; Huettig, 2015; Molinaro et al., 2016; Pickering & Gambi, 2018; Pickering & Garrod, 2013). One such proposal (Pickering & Garrod, 2013) assumes that prediction is based on covert simulation: motor-to-sensory forward models used to monitor one’s own speech during production (Hickok et al., 2011; Tourville & Guenther, 2011) would also be implemented during comprehension, to predict forthcoming information.

A straightforward way to test whether predicting others’ speech relies on running a simulation through our own production machinery is offered by the investigation of prediction in people with impaired speech planning and execution. People who stutter (PWS)^1^ represent a suitable population for this aim. Developmental stuttering (DS) is a multifactorial disorder characterized by disruption of the normal speech flow, resulting in blocks, pauses, repetitions, and prolongations of speech sounds. Although multiple factors contribute to stuttering and the exact etiology remains elusive, core deficits have been linked to dysfunctions in neural structures devoted to the planning, timing, and execution of speech-motor sequences. Compared to typically developed speakers, PWS show aberrant morphology and function of cortical regions such as the primary sensorimotor areas, the supplementary motor area (SMA) and the pre-SMA, the inferior frontal gyrus, temporo-parietal areas, the basal ganglia and the cerebellum (Alm, 2004; Busan, 2020; Chang et al., 2019; Etchell et al., 2018). Investigations on white matter structure further support this picture, revealing that, with respect to controls, PWS show atypicalities in speech-motor pathways (Etchell et al., 2018; Kronfeld-Duenias et al., 2016a, 2016b, 2018; Neef et al., 2021).

Neurocomputational models of speech production, such as the DIVA/GODIVA (Bohland et al., 2010; Guenther, 2016; Tourville & Guenther, 2011) and the HSFC (Hickok, 2012; Hickok et al., 2011) models, provided theoretical ground for the interpretation of these findings. Despite differing with respect to some architectural aspects, both frameworks stress the relevance of sensorimotor integration for planning and producing speech (for a discussion and comparison, see Guenther & Hickok, 2015). In both frameworks, motor plans are employed to predict auditory (and somatosensory) consequences, and these pieces of information are integrated in order to obtain efficient speech production. In this context, DS has been proposed to be caused by impaired feedforward processing of speech-motor plans (Chang & Guenther, 2020; Civier et al., 2013), as well as by inefficient motor-to-sensory predictions (Hickok et al., 2011; Max et al., 2004) and/or by an overreliance on the feedback control system (again, as a consequence of an inefficient and more inhibited feedforward system) (Civier et al., 2010; Max et al., 2004).

The investigation of speech planning with techniques that enable a high temporal resolution provided compelling evidence in support of this picture. In particular, by decomposing the electro-/magnetoencephalographic signal, it is possible to observe how the power of different frequency bands is modulated over time as a function of cognitive, sensory, and motor processing. Of particular interest in this context are the alpha (8-12 Hz) and beta (13-30 Hz) frequency bands. In people without speech production deficits, a power decrease in these frequency bands is found during speech planning and execution. This power decrease is thought to reflect the engagement of regions associated with memory and motor processes for language and speech production (Piai & Zheng, 2019; Saltuklaroglu et al., 2018). In the case of DS, these frequencies have been shown to be abnormally modulated – especially in motor and premotor associative regions – in syllable and word production tasks, indexing inefficient motor-to-sensory transformation and reduced coordination in engaging cortical regions devoted to speech production (Jenson et al., 2018; Joos et al., 2014; Mersov et al., 2016; Mock et al., 2016).

Given impaired speech-motor control in DS, if speech planning and motor-to-sensory forward modeling contribute to prediction during speech comprehension (Garrod et al., 2014; Pickering & Garrod, 2013), then it can be suggested that PWS would show impaired/atypical prediction processes during comprehension. In the present study we tested this hypothesis. To this end, we focused on two electrophysiological correlates of predictability, namely the pre-target alpha‒beta power decrease and the post-target N400 event-related potential (ERP) effect.

In non-clinical populations, alpha and beta power decreases before predictable with respect to unpredictable (but still plausible) words have been observed during written and auditory language comprehension (Armeni et al., 2019; Gastaldon et al., 2020; Molinaro & Monsalve, 2018; Rommers et al., 2017; Wang et al., 2018). Given the modulation in the same direction of the same frequency bands for language production, some researchers suggested that these power decreases may underlie at least partially common processes for predicting and producing language (Molinaro et al., 2016). A recent study investigating oscillatory modulations in auditory language prediction and in context-induced word production provided the first clear evidence in such direction. Gastaldon et al. (2020) showed that the pre-target power modulations associated to predictability in the two tasks are positively correlated in left-lateralized language areas, with stronger correlations in frontal regions, more prominently involved in spoken word production. Critically, these frontal regions are implicated in impaired speech planning and inefficient timing of speech sequence initiation in DS (Etchell et al., 2018). To the best of our knowledge, no study has investigated oscillatory modulations in language comprehension in PWS. The present study will fill this gap in the literature. Instead, differences between fluent speakers and PWS have been reported in ERP studies investigating the N400 response to semantic violations during comprehension. Typically, anomalous words embedded in a sentence elicit a stronger negative-going deflection relative to regular words in centro-parietal electrodes after 300-500 ms from word presentation. For instance, Murase et al. (2016) and Weber-Fox (2001) found a reduced N400 effect in response to written semantic violations in PWS relative to controls. The authors interpreted these results as an index of impaired semantic integration (but see Weber-Fox & Hampton, 2008, for a failure to detect an effect in the auditory modality). The N400 effect is also elicited when contrasting the same word embedded in neutral *vs*. constraining sentences (i.e., non-predictable *vs*. predictable words), making the N400 a strong EPR marker of facilitatory predictive processing (Hodapp & Rabovsky, 2021; Kuperberg et al., 2020; Kutas & Federmeier, 2011; Nicenboim et al., 2020; Nieuwland et al., 2020; Urbach et al., 2020). No previous study tested whether the N400 response in PWS is altered by employing non-anomalous non-predictable *vs*. predictable words. Providing such evidence is a further aim of the present study.

In the experiment, adults who stutter (AWS) and typically fluent adults (TFA) performed a sentence comprehension task, in which the predictability of the final word was manipulated by the preceding sentential context. Participants listened to a sentence frame and then, after a silent interval of 800 ms, they listened to the target word. Sentence frames could be highly (HC) o low constraining (LC), thus making the target predictable or non-predictable, respectively. Oscillatory activity has been analyzed in the silent interval between the sentence frame and the target. ERPs have been computed at the onset of the target word, analyzing the N400 time-window (300-500 ms post-target) (see Figure 1). If DS affects the efficiency of predicting during auditory language comprehension, we expected AWS to show a reduced alpha‒beta power decrease before the critical target word and a reduced N400 effect, relative to TFA.

**Figure 1.**
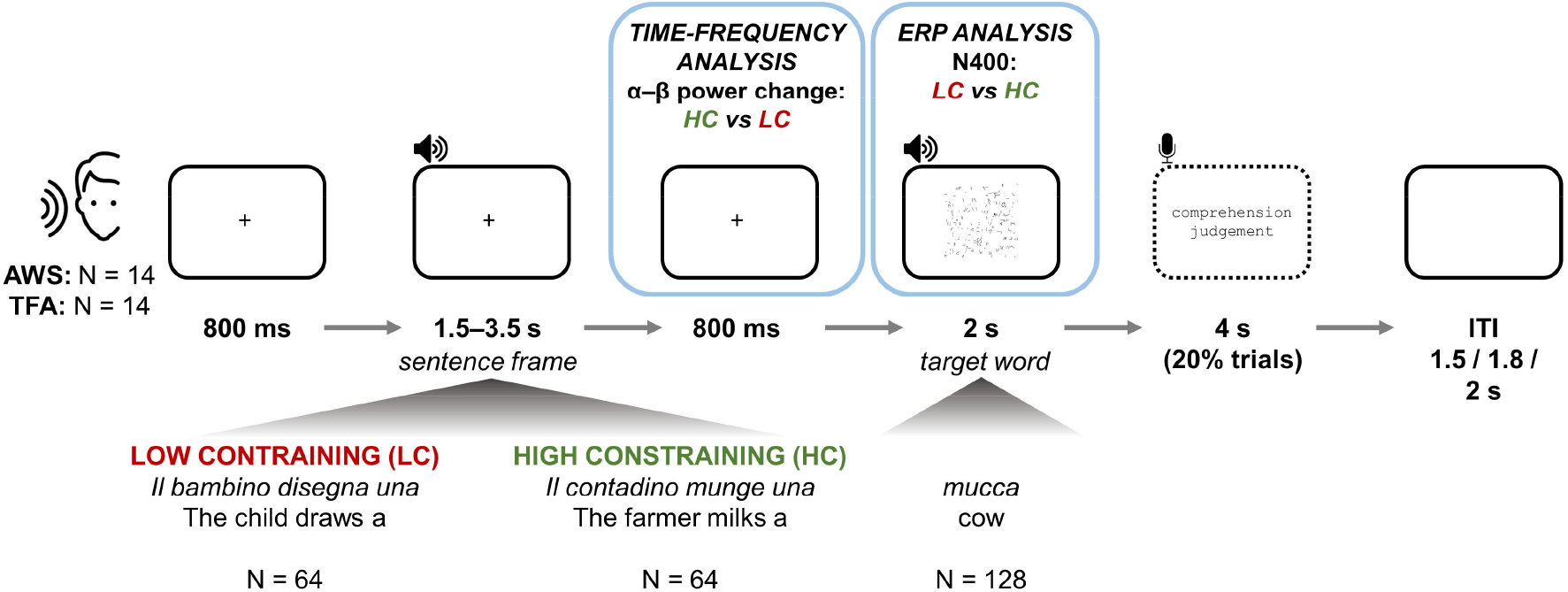
Task structure and analyses of interest.

## Materials and Methods

### Participants

Sample size was not determined prior to data collection. Participant recruitment in stuttering research, as with other special populations for disorders at low prevalence, is inherently sub-optimal (Jones et al., 2002). Given the higher incidence of PDS in males (Smith & Weber, 2017), we chose to recruit males only in order to have a more homogeneous sample. This was also justified by research showing sex differences in neural responses, that may be the consequence of intrinsic sex-related characteristics, as well as the different effects of adaptation to PDS or the consolidation of fluency-inducing strategies (Busan et al., 2013; Fox et al., 2000). As a consequence, the final sample included 14 right-handed male adults who stutter (AWS), recruited on a voluntary basis (mean age = 33.29 years, sd = 9.06; handedness evaluated by means of the Edinburgh Handedness Questionnaire, Oldfield, 1971; mean laterality index = 81.07, sd = 22.38). To be included in the study, AWS either had been diagnosed from a speech pathologist or clearly exhibited overt stuttered speech, with reported idiopathic stuttering starting during childhood. For the control group (typically fluent adults, TFA), the data for 9 male participants collected for Gastaldon et al. (2020) was pooled with newly collected data from 5 male participants recruited on a voluntary basis, for a total of 14 age- and handedness-matched male adult controls (mean age = 31.79 years, sd = 9.22; mean laterality index = 83.93, sd = 13.33). These participants were all fluent speakers, with no history of DS or other developmental disorders. All participants were native speakers of Italian. The two groups did not significantly differ for age (t = 0.434, df = 25.992, p = 0.668) or handedness (t = -0.411, df = 21.192, p = 0.686). History of psychiatric or neurological disorders, diagnosis of language or cognitive deficits, and ongoing or recent use of medications that could influence the functioning of the central nervous system were part of the exclusion criteria for both groups. All participants signed an informed consent prior to the session and were offered a compensation of 15€ for taking part to the study. The study was conducted in accordance with the Declaration of Helsinki and was approved by the Ethics Committee for Psychological Research of the University of Padova (protocol n. 3073).

### Stimuli

The stimuli are the same as employed in Gastaldon et al. (2020). One hundred twentyeight nouns associated to concrete concepts were selected and associated to corresponding black-and-white line pictures (size: 240 × 240 pixels). A scrambled version was generated for each picture, so that the referent was not recognizable. Each noun was associated to two sentence frames, one whose semantic content allows to easily predict the target word (high constrain, HC; cloze probability mean = 0.873, sd = 0.092) and one for which it is not possible to predict the target, despite still being a plausible conclusion to the sentence frame (low constraint, LC; cloze probability mean = 0.052, sd = 0.078). The total set resulted in 256 sentence frames (128 HC, 128 LC). The sentence frames for each target word were matched for number of syllables, were kept similar in their syntactic structure, and had the same final word. Sentence frames and target words were recorded separately by a female native speaker in a quiet room using a microphone connected to a PC and using the software Audacity (sampling rate of 44.1 KHz). The speaker was instructed to keep the reading pace as steady as possible and to keep a constant distance from the microphone. Recordings were then appropriately trimmed at the beginning and at the end using Audacity. The approximate number of syllables per second for each sentence frame, assuming a constant pace, was estimated as the number of syllables of the sentence divided by the length of each audio file. This measure was used to ensure a similar pacing across conditions. Target words and their associated sentence frames were then divided into two lists, A and B, each containing 64 target words and the associated 128 sentence frames and matched for a number of variables. Importantly, there were no significant differences of CP values within HC and LC conditions between the two lists (i.e., they elicited the same level of constraint) (please, refer to the Supplementary Material for all aspects of stimuli matching).

### Stuttering severity evaluation

The Stuttering Severity Instrument-4 (SSI-4; Riley, 2009) was used to evaluate stuttering severity in AWS. Before the experiment, participants were asked to 1) read a brief passage (394 syllables) and 2) spontaneously talk about a topic of their choice (hobby, school, work etc.). Participants were audio- and video-recorded during this session. The recordings were then inspected by a psychologist with experience in stuttering evaluation (G.N.) and checked by one of the authors (P.B.), to evaluate the percentage of stuttered syllables, the longest block durations, and face and limb movements associated with dysfluencies, in both tasks. Subsequently, a total score was assigned to each participant (SSI-4 score henceforth).

### Procedure

Participants sit in a soundproof room. Stimuli were delivered with E-Prime 2.0 (Psychology Software Tools, Pittsburgh, PA) on a computer with a CRT monitor and built-in speakers. The same paradigm was used in two sessions, one with a comprehension task and one with a production task. After listening to the sentence frame, in the comprehension task participants listened to the final target word, while in the production task they named a picture appearing on the screen to complete the sentence frame (see Gastaldon et al., 2020). Participants performed the two tasks in separate blocks. Task-specific instructions were given before each block. For each participant, the two stimuli lists (A and B) were assigned to the tasks, resulting in a 2×2 design. Task order and list-task association were counterbalanced across participants. Here we focus on the comprehension task (see Figure 1); the results of the production task are reported in the Supplementary Material.

After a fixation cross (800 ms), the auditory sentence frame was presented, while still displaying a fixation cross. At the end of the frame, a silent gap of 800 ms was introduced before the auditory target word was presented (concomitantly with a scrambled picture of the referent). The scrambled picture was introduced to keep consistency with the production task (i.e., to have a visual stimulus in both tasks; see Supplementary Material). To ensure that participants engaged in the task, 20% of the trials included a true/false judgement about the sentence, presented as a written statement on the screen after the target. Participants were asked to answer true/false by means of vocal response, recorded by means of a microphone positioned at approximately 50 cm from the participant. Inter-trial interval consisted in a blank screen presented at varying intervals (1.5, 1.8 and 2 seconds). Trial order was pseudo-randomized for each participant by using Mix (van Casteren & Davis, 2006). The same condition was not repeated for more than three times consecutively, and between the first and the second presentation of the same target there were at least 7 trials. Every 32 trials, participants could take a pause. A familiarization phase was carried out before the task (8 trials, no stimuli were presented in the subsequent experiment). The task lasted approximately 20 minutes.

### EEG recording and pre-processing

Electroencephalogram (EEG) was recorded with 64 active Ag/AgCl electrodes system (ActiCap, Brain Products) with a 10-20 placement convention. Sixty electrodes were used as active electrodes (Fp1, Fp2, AF3, AF4, AF7, AF8, F1, F2, F3, F4, F5, F6, F7, F8, Fz, FT7, FT8, F1, F2, F3, F4, F5, F6, Fz, FC1, FC2, FC3, FC4, FC5, FC6, T7, T8, C1, C2, C3, C4, C5, C6, Cz, TP7, TP8, CP, CP2, CP3, CP4, CP5, CP6, CPz, P1, P2, P3, P4, P5, P6, P7, P8, PO3, PO4, PO7, PO8, PO9, PO10 POz, O1, O2, Oz), while 3 were used to record blinks and saccades (external canthi and below the left eye). Reference was placed at the left earlobe. Impedance was kept below 10 kΩ throughout the experiment. The signal was amplified and digitized at a sampling rate of 1000 Hz. Each task was recorded separately. All pre-processing steps were performed using the MATLAB toolbox Brainstorm (Tadel et al., 2011). A high-pass filter at 0.5 Hz with 60 dB attenuation was applied to the raw data. Noisy or flat channels were excluded (max 2 channels per participant). No interpolation of excluded channels was performed. Segments with extreme muscle artifacts were excluded. Subsequently, Independent Component Analyses (ICA) with 60 components was computed on the continuous recordings to detect and remove artifact components with known time-series and topographies (blinks, saccades, and power-line noise at 50 Hz). If any channels were removed from the analyses due to excessive noise, the number of components for the ICA was reduced to the number of available channels. Markers for incorrect responses were manually added to the continuous EEG recording according to the off-line evaluation of the audio files. Epochs were imported around three event-markers: (a) the onset of the first fixation cross (from -1.5 to 1.5 s), and (b) the onset of the 800 ms gap between sentence frame and target (from -1.5 to 1.5 s), and (c) the onset of the auditory target word (from -0.3 to 1 s, baseline corrected in the inverval -300 to 0 ms). Epochs (a-b) were employed for time-frequency analyses, while (c) for ERP analyses. While epochs (b) were divided into HC and LC conditions, epochs (a) were used as a condition-average baseline for the normalization of epochs (b). All epochs were visually inspected, and those with artifacts (uncorrected blinks/saccades, muscle activity, channel drifts, transient electrode displacements) were rejected. Additionally, trials that were associated to an incorrect response to the true/false judgement were discarded from the analyses. The mean percentage of epochs retained for each group are the following: AWS: baseline: 91.9%%, interval_HC_: 91%, interval_LC_: 89.7%, target_HC_: 88.8%, target_LC_: 86.4%; TFA: baseline: 88.8%%, interval_HC_: 90%, interval_LC_: 87.7%, target_HC_: 89.5%, target_LC_: 87.5%

### Source estimation and ROI definition

To perform source analysis, the head model was built using OpenMEEG BEM (Boundary Element Method) with 8002 vertices as forward solution (Gramfort et al., 2010) and ICBM152 as template anatomy. A noise covariance matrix, providing a specification of channel noise, was computed by taking the baseline epochs in the time-window [-550 - 250] ms. The head model and the noise covariance matrix were used for the Minimum Norm Imaging (NMI) with the sLORETA (Standardized Low Resolution Brain Electromagnetic Tomography; Pascual-Marqui et al., 1994) as inverse solution. Given the lack of individual anatomy scans, the dipole orientation was set to be unconstrained.

Subsequently, we defined regions of interest (ROIs) by generating a custom atlas starting from the Desikan-Killiany atlas (Desikan et al., 2006) as implemented in Brainstorm (see Table S1 and Figure S1 for ROI definition and the projection on the cortex). ROIs were defined by taking into consideration the literature on the neural bases of developmental stuttering and on linguistic processing (see references in the Introduction). Time-series at the ROI level were extracted by using Principal Component Analysis (PCA) and selecting the first component, as implemented in Brainstorm. These time-series were then the input for the time-frequency decomposition.

### Time-frequency decomposition and analyses

Time-frequency decomposition was performed both at the sensor- and the ROI source-level by using the Morlet wavelet implementation in Brainstorm. The decomposition was performed for each trial in the range 8-30 Hz, with a linear step of 1 Hz, by generating wavelets from a mother wavelet with central frequency = 1 and FWHM = 3. After obtaining the average time-frequency maps of each condition (baseline, HC, and LC) for each participant, the HC and LC conditions were normalized against the baseline, specifically in the pre-trial interval from -550 to -250 ms. The normalization method was the event-related synchronization/desynchronization (ERSD), which yields the %-power change relative to the baseline.

Due to the small sample size and to the contrasts between groups leading to lower statistical power, we performed both corrected (cluster-based permutations, Maris & Oostenveld, 2007; number of Monte Carlo simulations = 1000) and uncorrected t-tests. All contrasts for time-frequency power modulations focused on the 800 ms silent time-window between the sentence frame and the target. When contrasting the HC and LC conditions within each group, one-tailed paired t-tests were performed (H0: HC ≥ LC, H1: HC < LC; α-level = 0.05). This choice was guided by previous literature that consistently showed a desynchronization in the alpha and beta bands pre-target associated to predictability (Gastaldon et al., 2020; Molinaro & Monsalve, 2018; Rommers et al., 2017; Wang et al., 2018). In this way, the research hypothesis is appropriately translated into a statistical hypothesis (Cho & Abe, 2013). To compare the predictability effects between groups in the two tasks and in absence of no previous studies contrasting the two populations, we contrasted the effects (i.e., the differences between conditions, Δ%power-change from now on) between AWS and TFA by means of independent two-tailed t-tests (H0: HC = LC, H1: HC ≠ LC; α-level = 0.05). When testing at the sensor-level, the minimum number of neighboring channels was set to 2, whereas when testing at the ROI level, the parameter was set to 0. This is due to the fact that, while at sensor level the activity of one sensor is affected by the activity from neighboring sensors, the components identified by the PCA at source level should be treated separately.

### Event-realated potential analyses

After pre-processing, all retained epochs were averaged separately by condition, and a low-pass filter at 40 Hz (60 dB attenuation) was applied to the average. Cluster-based permutations on the 300‒500 ms interval typical of the N400 component were performed to test the difference between non-predictable and predictable words, corresponding to LC and HC sentence frames respectively, within each group (paired one-tailed t-tests; H_0_: LC ≥ HC, H_1_: LC < HC; α-level = 0.05; number of Monte Carlo simulations = 2000; minimum number of neighboring channels = 2). A cluster-based permutation was performed on the differentials (non-predictable – predictable, ‘N400’ for simplicity) between the AWS and the TFA group on the same time-window to test for an interaction. Given our research hypothesis and results from previous research, we employed an independent one-tailed t-test (H_0_: N400_AWS_ ≤ N400_TFA_, H_1_: N400_AWS_ > N400_TFA_; α-level = 0.05; 2000 Monte Carlo simulations; minimum number of neighboring channels = 2).

### Correlations

Pearson’s correlations were performed to better understand the relation between power modulations, ERP responses, and behavioral measures. Power changes in each ROI were averaged across the 800 ms pre-target time-window in a priori frequency ranges (alpha: 8-12 Hz; beta1: 12-20 Hz; beta3: 20-30 Hz), and correlated with the mean amplitude of the N400 effect (calculated in the 300-500 ms time-window across centro-parietal sensors where the N400 effect is typically evident (CP3, CP1, CPz, CP2, CP4, P3, P1, Pz, P2, P4, PO3, POz, PO4), and, for the AWS group only, with the SSI-4 values.

## Results

Statistical contrasts of time-frequency data within (Δ = HC *vs*. LC) and between groups (Δ_AWS_ *vs*. Δ_TFA_) at the sensor level are shown in Figure 2. For the source-level, between group contrasts are shown in Figure 3 (for sake of simplicity, within-group contrasts are reported in Figures S2 and S3). As specified in the Methods section, we performed both cluster-corrected and uncorrected t-tests; only the latter analysis yielded statistically significant results.

**Figure 2.**
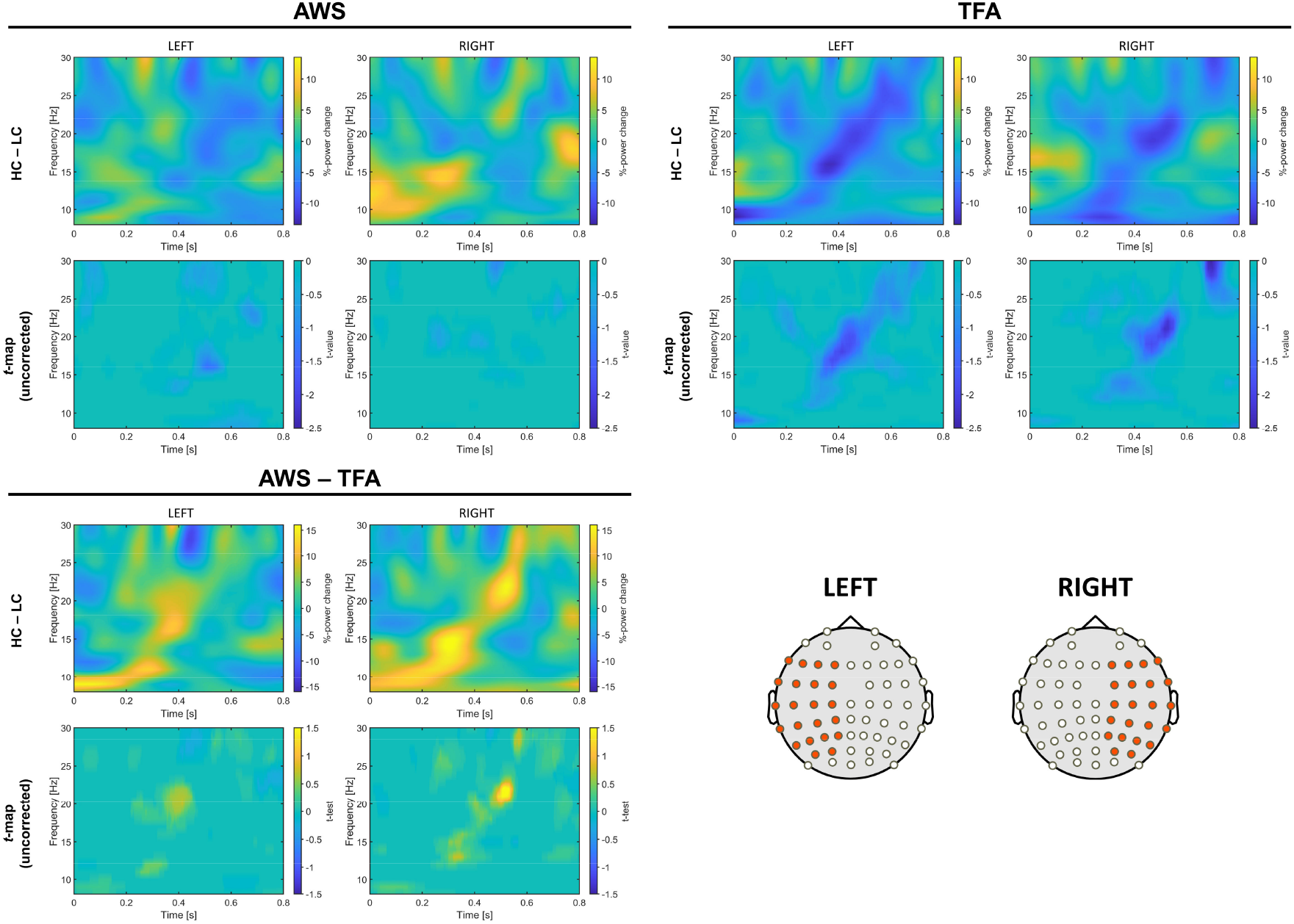
Time-frequency results at the sensor level. Results are averaged across the sensors in the two hemispheres as indicated on the scalp models in the bottom-right panel (only for visualization purposes). For each panel (AWS, TFA, and AWS ‒ TFA), the top row shows the Δ%-power change in the difference between HC and LC conditions, while the bottom row represents the t-values of uncorrected statistical tests (p < 0.05).

**Figure 3.**
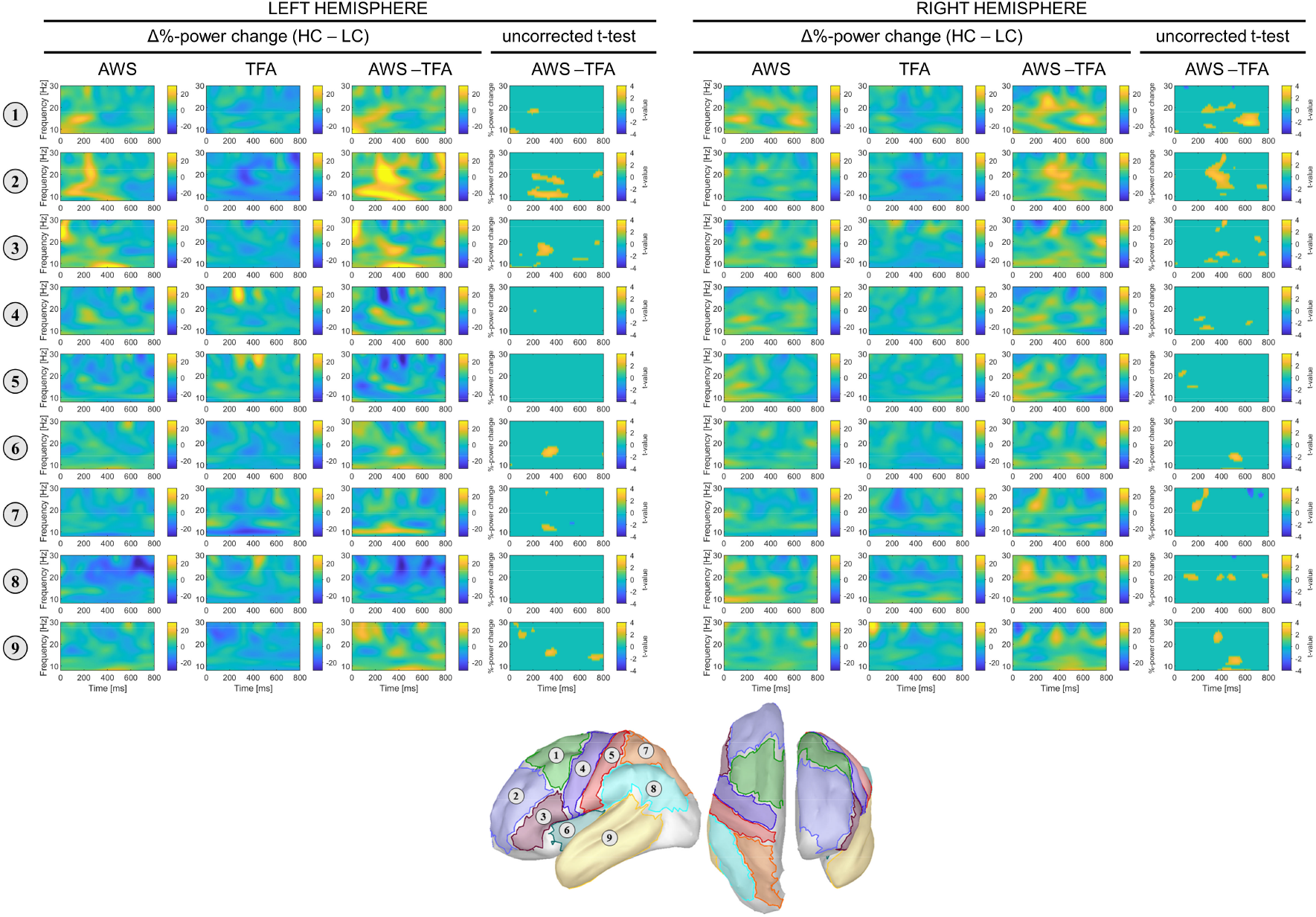
Time-frequency results at the source ROI level. For each hemisphere, Δ%-power change for AWS, TFA, and AWS – TFA, and the map of uncorrected t-tests for AWS *vs*. TFA (p < 0.05) are shown. Each numbered row corresponds to a ROI depicted on the cortical model below. For the contrasts within each group, please see Figures S2 and S3.

At the sensor level, while TFA display a clear power decrease before predictable (HC) relative to non-predictable (LC) words, especially in the beta range, this effect is less marked in AWS. Contrasts between AWS and TFA revealed more prominently positive than negative t-values, especially in the beta range (13-25 Hz). This indicates that the predictability effect elicits lower power in TFA relative to AWS.

This pattern is confirmed at the source level (Figure 3). Power decrease is visibly marked in prefrontal, frontal, temporal, and parietal regions, bilaterally, in TFA. In AWS, the effect seems to be limited to posterior (temporal and parietal) regions of the left hemisphere. Statistical contrasts between the effects in AWS and TFA revealed positive t-values throughout the considered ROIs. More specifically, this difference results from a less marked power decrease in posterior regions in AWS relative to TFA, but from a power increase in anterior regions in AWS, whereas TFA consistently show a power decrease.

The cluster-corrected statistical analyses on the ERPs (Figure 4) revealed that non-predictable *vs*. predictable words elicited the well-known N400 effect in both groups (TFA: p = 0.0005, t-sum = -32112, size = 8806; AWS: p = 0.001, t-sum = -19210, size = 6799). However, the contrast between groups revealed that the N400 effect is reduced in AWS, reflected in less negative values (p = 0.032, t-sum = 3516, size = 1408).

**Figure 4.**
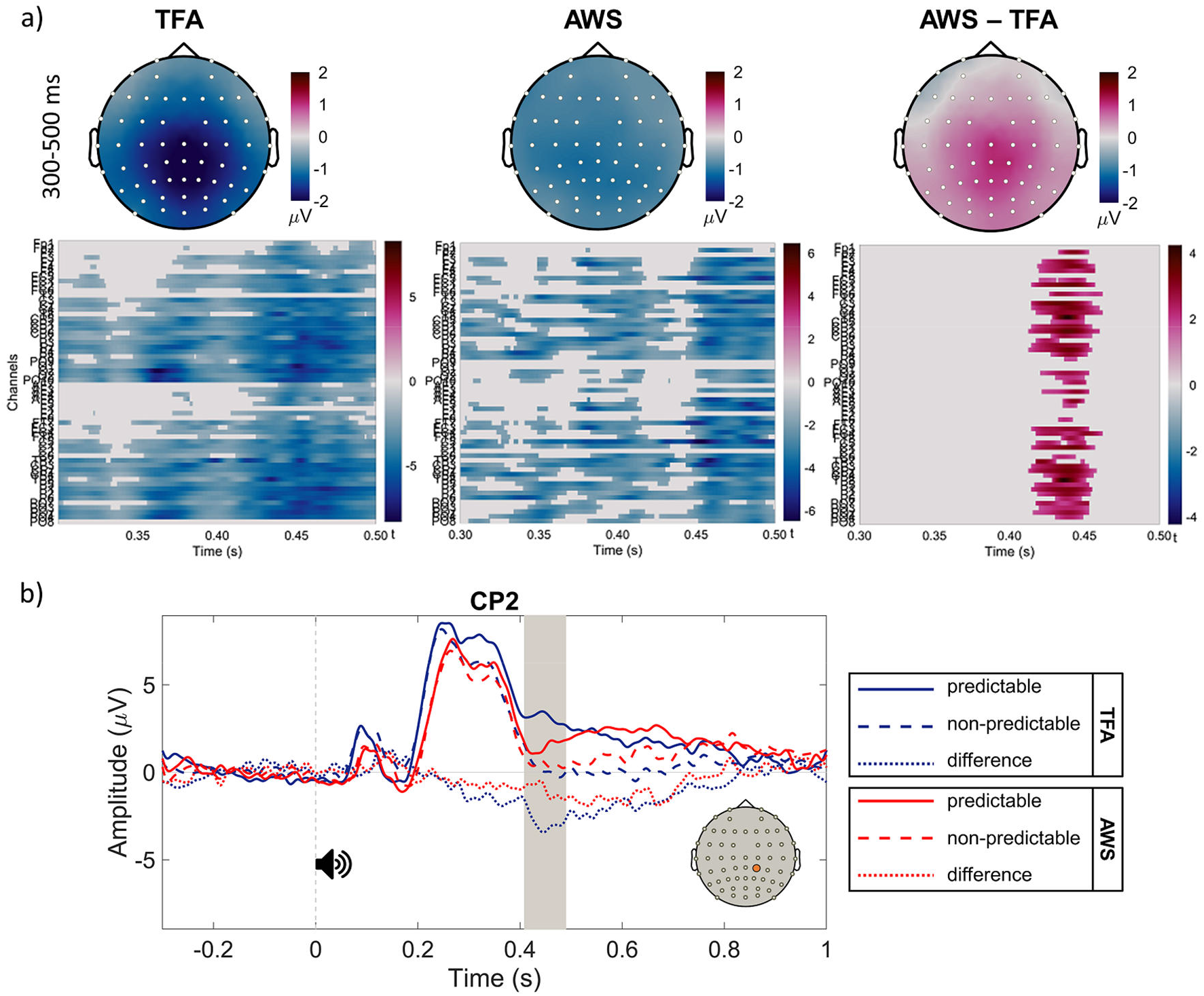
N400 effects within and between groups. Panel (a): Topographical plots of the voltage difference between non-predictable and predictable targets averaged in the 300-500 ms post-target time-window for the TFA group (left), the AWS group (center), and the difference between AWS and TFA groups (right). The raster plots below show the significant clusters identified. Panel (b): Grand-average waveforms from a representative electrode, CP2 (highlighted in orange on the scalp model). Time 0 signals the onset of the auditory target word. The dotted lines represent the N400 effect for AWS (in red) and TFA (in blue). The area in grey highlights the time interval where the significant cluster between groups is defined.

We further explored the relationship between DS and prediction by performing correlations between pre-target power change in regions of interest and i) stuttering severity (SSI-4 scores), and ii) post-target N400 amplitude. Correlations between power modulations and N400 revealed that, in AWS, the magnitude of the N400 effect is positively correlated with pre-target alpha (8-12 Hz) power modulation in premotor regions of the right hemisphere, comprising the supplementary motor regions (R = 0.61, p = 0.022): less negative values (a weaker effect) are associated to a lower power decrease or a power increase in this region. No significant correlations were found with stuttering severity. In TFA, the N400 magnitude is positively correlated with beta1 (13-20 Hz) (R = 0.58, p = 0.029) and beta2 (20-30 Hz) (R = 0.54, p = 0.048) power changes in the left inferior frontal cortex: a stronger effect is associated to a stronger power decrease. These correlations are shown in Figure 5.

**Figure 5.**
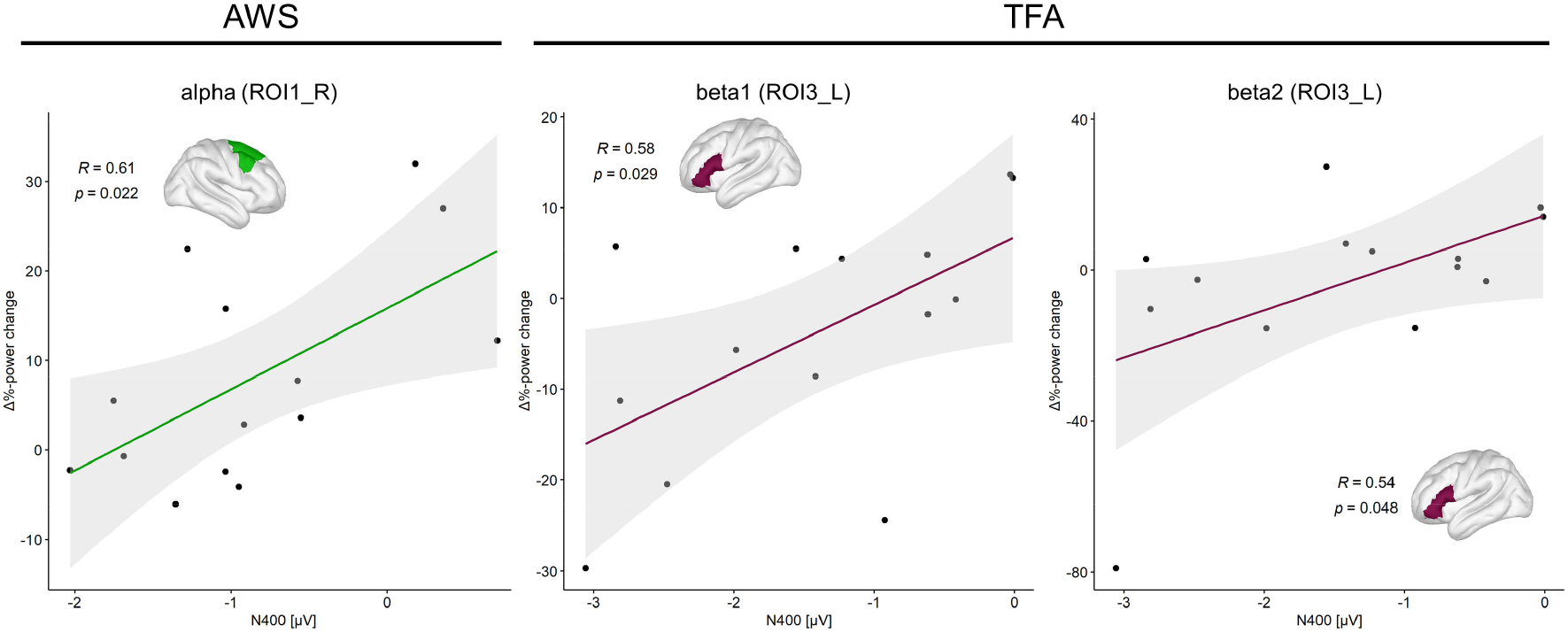
Correlations between ROI Δ%-power change and the N400 effect in the two groups. Alpha range: 8-12 Hz; beta1 range: 13-20 Hz; beta2 range: 20-30 Hz.

## Discussion

In this study we contrasted electrophysiological correlates of predictability (pre-target alpha‒beta power decrease and post-target N400 effect) between a group of adults who stutter and a group of matched fluent controls, to test whether aspects of speech-motor control ‒ which are extensively defective in AWS ‒ play a role in prediction during spoken language comprehension. We found that in AWS the alpha‒beta power decrease usually found before predictable words is less consistent and primarily restricted to posterior areas. Convergently, the N400 effect usually reported for non-predictable *vs*. predictable words is significantly reduced in AWS. Significant correlations between power modulations and the N400 were also found, revealing that, in AWS, the amplitude of the N400 effect is positively correlated with alpha power modulations in right supplementary and associative premotor areas.

### Atypical alpha-beta power modulations and a reduced N400 effect signal inefficient prediction during spoken language comprehension in AWS

The power decrease in the oscillatory activity found when contrasting high *vs*. low constraining conditions was less marked in AWS than in the TFA group. At the source level, in AWS the power decrease was mainly evident in the low and high beta bands in left posterior areas. In TFA, the effects involved also anterior areas. When comparing differentials between AWS and TFA (i.e., when comparing the effects of predictability), differences were evident across many anterior and posterior regions in both hemispheres. This likely indicate that the two groups engaged in prediction in a different manner. Notably, group differences in the posterior areas were due to a reduced power decrease associated to predictability in AWS, whereas in frontal areas the differences were characterized by the presence of a power increase in stuttering. However, given the specific directional hypothesis formulated on the basis of previous findings, in the contrast between HC and LC conditions, performed within each group, we only tested the negative tail of the distribution. For this reason, the power increase observed in frontal areas in AWS may not be statistically significant. From a functional point of view, alpha‒beta power increase has been interpreted as a marker of cortical inhibition and maintenance of the current motor or cognitive state, while a power decrease has been suggested as a marker of cortical engagement, change in the current set, and it has been shown to be negatively related to the richness and precision of information retrieval and encoding, i.e., the stronger the decrease, the richer the information (Engel & Fries, 2010; Griffiths et al., 2019; Hanslmayr et al., 2012). Consequently, in the context of language prediction, the alpha‒beta power decrease has been linked to the pre-activation of representations, i.e., the access of information from long-term memory before the stimulus is actually encountered and processed (Prystauka & Lewis, 2019). The pattern emerging from our results suggests that i) the information pre-activated by AWS when predicting is less precise, or AWS are less committed to their predictions, and that ii) AWS may be also exerting an inhibitory control at some level of processing that engages frontal areas. Speculatively, this inhibitory control may be carried out in order to avoid an “incorrect” activation of information that may not be strictly related to the predicted target, thus inhibiting possible competing information. In fact, AWS have been found to show an altered inhibitory control in tasks such as motor sequences (Busan et al., 2020), but also during lexical selection for word production (Maxfield, 2020). This suggests the existence of altered signal-to-noise ratios for action release in DS, thus resulting in impaired inhibitory processing (compare with Alm, 2004). The present findings are compatible with this recent evidence, suggesting that altered inhibition processes may play an important role in sensorimotor aspects of DS (see Neef et al., 2016, 2018), impacting also aspects of language comprehension, such as prediction.

As for the results obtained in the ERP analyses, even though both groups showed a N400 effect (i.e., the difference between non-predictable and predictable targets), the effect was significantly attenuated in AWS. Importantly, the responses to non-predictable targets are comparable between the two groups and the difference is driven by the ERP response to predictable targets, which resulted in a more positive-going deflection in TFA than in AWS. This strongly suggests that the difference in the N400 effect reflects a specific difference in how predictable words are processed, rather than a general difference in word processing. The different amplitude observed only for predictable words could reflect the reduced top-down facilitation of bottom-up processing, due to less efficient predictions. Arguably, a malfunctioning speech-motor system is unable to appropriately predict the auditory consequences of speech-motor plans; predicting during comprehension by means of the same mechanisms brings about inefficient auditory (phonological) prediction of others’ speech.

Interestingly, a positive correlation between power modulation and the magnitude of the N400 effect in AWS showed that higher alpha power in posterior dorsal frontal regions (ROI 1) ‒ which implies a reduced cortical engagement ‒ is associated to a reduced N400 effect (less negative voltage). Such ROI includes the premotor cortex, the SMA and pre-SMA. These regions are anatomically and functionally connected with both subcortical (basal ganglia, thalamus) and cortical structures (anterior cingulate cortex, inferior frontal gyrus, angular gyrus, pre-motor cortex, primary motor cortex, somatosensory cortex and middle cingulate cortex) (Hertrich et al., 2016). These connections make premotor regions and the SMA complex important nodes in the extended speech-motor network (Alario et al., 2006; Ghosh et al., 2008). In the DIVA/GODIVA framework of speech planning and execution, the SMA and pre-SMA are responsible for the sequencing and timing of speech-motor units (Guenther, 2016). Recent research has shown that these mechanisms play a role in auditory perception and imagery of speech, by mediating the generation of top-down auditory predictions (Lima et al., 2016). Importantly, the SMA complex (together with regions such as the inferior frontal cortex, especially in the left hemisphere) represents a crucial node in the pathophysiology of DS: dysfunction in the cortico-basal-thalamo-cortical system – comprising the SMA complex – causes a disruption in the internal timing of the initiation of planned, complex motor sequences, such as those required for speech production, leading to stuttering (Busan, 2020; Busan et al., 2019; Chang & Guenther, 2020). Compatibly, premotor and SMA regions of the right hemisphere have been reported as being part of a plausible compensation system, that may be useful to overcome dysfluencies in stuttering (Brown et al., 2005; Busan et al., 2019; Neef et al., 2015, 2016, 2018). Considering all these lines of evidence, we suggest that during prediction the SMA-complex may be (bilaterally) involved in the coordination of sensorimotor information flow between cortical areas. The positive correlation between cortical inhibition in this region (reflected in alpha power increase) and the reduced efficiency in word prediction (reflected in a reduced magnitude of the N400 effect) supports the hypothesis that a network involved in motor-auditory mapping and sequence processing is relevant for prediction during spoken language comprehension, and that a disfunction in such network negatively affects prediction efficiency. For what concerns correlations in the control group, they show that a power decrease in the beta range in the left inferior frontal cortex (i.e., a stronger engagement of this cortical region) is associated to a stronger N400 effect (more negative values). The involvement of this region is in line with previous studies (Jakuszeit et al., 2013; Lau et al., 2008; Siman-Tov et al., 2019; Wang et al., 2018). Interestingly, the inferior frontal cortex has functional and anatomical connections with the SMA complex. If these connections are dysfunctional in DS, as shown in the literature, the information flow may be disrupted, resulting in compromised predictions. While the correlation between power modulations and the N400 effect is found in the left hemisphere in TFA, for AWS the correlation involves the right hemisphere. As noted above, the recruitment of (especially motor and premotor) regions in the right hemisphere is often reported as a compensation mechanism; for this reason, the different laterality of the correlations found in the two groups is not surprising.

The present findings add also relevant evidence in favor of the strict interdependence of brain areas and neural processes that support speech production and self-monitoring, as hypothesized by the DIVA/GODIVA model (Guenther, 2016). Within the model, and in a very simplified manner, while frontal regions may be involved in evaluating the current context for the generation of appropriate predictions, this information may be promptly delivered to the motor system, for its possible initiation (e.g., the SMA complex). Similarly, information is sent to temporal and parietal regions for auditory and somatosensory mapping and prediction. This will allow the system to be “ready”, both if some corrections are necessary (with respect to the perceptual information received), or if it will be needed to respond with a prompt action (i.e., speech). Within such dynamic and integrated system, a sub-optimal feedforward performance of the sensorimotor system of AWS may easily result in inefficient planning and prediction skills (see also Chang & Guenther, 2020 and Civier et al., 2013 in the context of the relations between stuttering and the DIVA/GODIVA model). In turn, this lower efficacy may trigger a cascade of sub-optimal responses, influencing also sensorial maps, and finally resulting in a reduced efficiency in proactive processing during speech comprehension.

### Limitations

The study also presents some limitations. First, the number of AWS we were able to recruit was quite restricted. This is due to the fact that participant recruitment in stuttering research, as with other special populations with disorders at low prevalence, is inherently sub-optimal (Jones et al., 2002). A second threat to generalizability is our choice of investigating only males. This choice was motivated by i) the higher incidence of PDS in males (Smith & Weber, 2017), and ii) attested biological sex differences in neural responses in AWS, that may be the consequence of intrinsic sex-related characteristics, as well as the different effects of adaptation to PDS or the consolidation of fluency-inducing strategies (Busan et al., 2013; Fox et al., 2000). Finally, possibly related to the previous points, we recognize the fact that, from a strict statistical point of view, the effects we obtained were significant only without correcting for multiple comparisons. Given the highly explorative nature of the study that aimed at exploring a likely subtle phenomenon in a clinical population, we believe that a less stringent approach is justifiable.

## Conclusions

We studied the electrophysiological correlates of prediction during auditory language comprehension in adults who stutter. We found evidence for less efficient prediction in this population. This study adds novel evidence in support of the involvement of the neural infrastructure devoted to speech-motor control in prediction during speech comprehension. The study ultimately stresses two major points: i) a more integrated investigation of language comprehension and production is desirable, and we advocate for a view in which the two modalities are seen as task sets drawing from at least partially shared resources, rather than two separate systems (see e.g., McQueen and Meyer, 2019), and ii) the investigation of on-line speech/language comprehension in DS, in addition to speech/language production, should be further pursued, in order to attain a better understanding of the neural architecture of language in healthy and/or pathological conditions. Specifically, regarding DS, we suggest that the role of executive functions and inhibitory control should be further investigated and taken as potential targets for better speech therapy interventions, extending treatment techniques using cognitive and linguistic approaches in addition to approaches mainly focused on “classical” fluency-shaping techniques. Future research should investigate which representations and processes are implicated in different tasks (and how), in order to better specify the language architecture and language use. This, in turn, can help in the development of more effective therapeutic approaches to developmental and acquired disorders of both language comprehension and production.

## Author contributions

S.G. and F.P. conceptualized the study. S.G. performed data collection and analyses. P.B. and G.A. provided methodological contributions for data analyses. S.G., F.P. and P.B. discussed and interpreted the results. S.G. drafted the manuscript, with contributions from F.P. and P.B. All authors reviewed the manuscript and approved the submission.

## Funding

S.G. was supported by a PhD fellowship from the University of Padova (2017-2020) and a research fellowship from the Dipartimento di Psicologia dello Sviluppo e della Socializzazione (DPSS), University of Padova (2020-2021). P.B. was supported by the Italian Ministry of Health (Project Code: GR-2018-12366027); the funding source was not involved in study design, in the collection, analysis and interpretation of data, in the writing of the report, and in the decision to submit the article for publication. G.A. was supported by the Italian Ministry of Health. We thank Elena Greatti for helping with data collection and Giulia Natarelli for evaluating the audio-video recordings and providing SSI-4 scoring.

*I dedicate this manuscript to the memory of my father Carlo (1946-2021). [S*.*G*.]

## Footnotes

1 In the literature, PWS is used to refer to both adults and children who stutter (AWS and CWS, respectively) indistinctively. The appropriate acronyms are used accordingly throughout the manuscript.

## Notes

### Competing Interest Statement

The authors have declared no competing interest.

## References

AbdulSabur, N. Y., Xu, Y., Liu, S., Chow, H. M., Baxter, M., Carson, J., & Braun, A. R. (2014). Neural correlates and network connectivity underlying narrative production and comprehension: A combined fMRI and PET study. Cortex, 57, 107–127. https://doi.org/10.1016/j.cortex.2014.01.017

Alario, F. X., Chainay, H., Lehericy, S., & Cohen, L. (2006). The role of the supplementary motor area (SMA) in word production. Brain Research, 1076(1), 129–143. https://doi.org/10.1016/j.brainres.2005.11.104

Alm, P. A. (2004). Stuttering and the basal ganglia circuits: A critical review of possible relations. Journal of Communication Disorders, 37(4), 325–369. https://doi.org/10.1016/j.jcomdis.2004.03.001

Armeni, K., Willems, R. M., van den Bosch, A., & Schoffelen, J.-M. (2019). Frequency-specific brain dynamics related to prediction during language comprehension. NeuroImage, 198, 283–295. https://doi.org/10.1016/J.NEUROIMAGE.2019.04.083

Bohland, J. W., Bullock, D., & Guenther, F. H. (2010). Neural representations and mechanisms for the performance of simple speech sequences. Journal of Cognitive Neuroscience, 22(7), 1504–1529. https://doi.org/10.1162/jocn.2009.21306

Brown, S., Ingham, R. J., Ingham, J. C., Laird, A. R., & Fox, P. T. (2005). Stuttered and fluent speech production: An ALE meta-analysis of functional neuroimaging studies. Human Brain Mapping, 25(1), 105–117. https://doi.org/10.1002/HBM.20140

Busan, P. (2020). Developmental stuttering and the role of the supplementary motor cortex. Journal of Fluency Disorders, 105763. https://doi.org/10.1016/j.jfludis.2020.105763

Busan, P., D’Ausilio, A., Borelli, M., Monti, F., Pelamatti, G., Pizzolato, G., & Fadiga, L. (2013). Motor excitability evaluation in developmental stuttering: A transcranial magnetic stimulation study. Cortex, 49(3), 781–792. https://doi.org/10.1016/j.cortex.2011.12.002

Busan, P., Del Ben, G., Russo, L. R., Bernardini, S., Natarelli, G., Arcara, G., Manganotti, P., & Battaglini, P. P. (2019). Stuttering as a matter of delay in neural activation: A combined TMS/EEG study. Clinical Neurophysiology, 130(1), 61–76. https://doi.org/10.1016/J.CLINPH.2018.10.005

Busan, P., Del Ben, G., Tantone, A., Halaj, L., Bernardini, S., Natarelli, G., Manganotti, P., & Battaglini, P. (2020). Effect of muscular activation on surrounding motor networks in developmental stuttering: A TMS study. Brain and Language, 205, 104774. https://doi.org/10.1016/j.bandl.2020.104774

Chang, S. E., Garnett, E. O., Etchell, A., & Chow, H. M. (2019). Functional and Neuroanatomical Bases of Developmental Stuttering: Current Insights. The Neuroscientist, 25(6), 566–582. https://doi.org/10.1177/1073858418803594

Chang, S., & Guenther, F. H. (2020). Involvement of the Cortico-Basal Ganglia-Thalamocortical Loop in Developmental Stuttering. In Frontiers in Psychology (Vol. 10, p. 28). Frontiers Media S.A. https://doi.org/10.3389/fpsyg.2019.03088

Chater, N., McCauley, S. M., & Christiansen, M. H. (2016). Language as skill: Intertwining comprehension and production. Journal of Memory and Language, 89, 244–254. https://doi.org/10.1016/j.jml.2015.11.004

Cho, H. C., & Abe, S. (2013). Is two-tailed testing for directional research hypotheses tests legitimate? Journal of Business Research, 66(9), 1261–1266. https://doi.org/10.1016/j.jbusres.2012.02.023

Christiansen, M. H., & Chater, N. (2016). The Now-or-Never bottleneck: A fundamental constraint on language. Behavioral and Brain Sciences, 39, e62. https://doi.org/10.1017/S0140525X1500031X

Civier, O., Bullock, D., Max, L., & Guenther, F. H. (2013). Computational modeling of stuttering caused by impairments in a basal ganglia thalamo-cortical circuit involved in syllable selection and initiation. Brain and Language, 126(3), 263–278. https://doi.org/10.1016/j.bandl.2013.05.016

Civier, O., Tasko, S. M., & Guenther, F. H. (2010). Overreliance on auditory feedback may lead to sound/syllable repetitions: Simulations of stuttering and fluency-inducing conditions with a neural model of speech production. Journal of Fluency Disorders, 35(3), 246–279. https://doi.org/10.1016/j.jfludis.2010.05.002

Clark, E. V, & Hecht, B. F. (1983). Comprehension, production, and language acquisition. Annual Review of Psychology, 34, 325–349.

Dell, G. S., & Chang, F. (2014). The P-chain: relating sentence production and its disorders to comprehension and acquisition. Philosophical Transactions of the Royal Society B: Biological Sciences, 369(1634), 20120394–20120394. https://doi.org/10.1098/rstb.2012.0394

Desikan, R. S., Ségonne, F., Fischl, B., Quinn, B. T., Dickerson, B. C., Blacker, D., Buckner, R. L., Dale, A. M., Maguire, R. P., Hyman, B. T., Albert, M. S., & Killiany, R. J. (2006). An automated labeling system for subdividing the human cerebral cortex on MRI scans into gyral based regions of interest. NeuroImage, 31(3), 968–980. https://doi.org/10.1016/j.neuroimage.2006.01.021

Engel, A. K., & Fries, P. (2010). Beta-band oscillations — signalling the status quo? Current Opinion in Neurobiology, 20(2), 156–165. https://doi.org/10.1016/J.CONB.2010.02.015

Etchell, A. C., Civier, O., Ballard, K. J., & Sowman, P. F. (2018). A systematic literature review of neuroimaging research on developmental stuttering between 1995 and 2016. Journal of Fluency Disorders, 55, 6–45. https://doi.org/10.1016/j.jfludis.2017.03.007

Fairs, A., Michelas, A., Dufour, S., & Strijkers, K. (2021). The Same Ultra-Rapid Parallel Brain Dynamics Underpin the Production and Perception of Speech. Cerebral Cortex Communications, 2(3), 1–17. https://doi.org/10.1093/TEXCOM/TGAB040

Fox, P. T., Ingham, R. J., Ingham, J. C., Zamarripa, F., Xiong, J.-H., & Lancaster, J. L. (2000). Brain correlates of stuttering and syllable production: A PET performance-correlation analysis. Brain, 123(10), 1985–2004. https://doi.org/10.1093/brain/123.10.1985

Gambi, C., & Pickering, M. J. (2017). Models Linking Production and Comprehension. In E. M. Fernández & H. S. Cairns (Eds.), The Handbook of Psycholinguistics (pp. 157–181). John Wiley & Sons, Inc. https://doi.org/10.1002/9781118829516.ch7

Garrod, S., Gambi, C., & Pickering, M. J. (2014). Prediction at all levels: Forward model predictions can enhance comprehension. Language, Cognition and Neuroscience, 29(1), 46–48. https://doi.org/10.1080/01690965.2013.852229

Gastaldon, S., Arcara, G., Navarrete, E., & Peressotti, F. (2020). Commonalities in alpha and beta neural desynchronizations during prediction in language comprehension and production. Cortex, 133, 328–345. https://doi.org/10.1016/j.cortex.2020.09.026

Ghosh, S. S., Tourville, J. A., & Guenther, F. H. (2008). A neuroimaging study of premotor lateralization and cerebellar involvement in the production of phonemes and syllables. Journal of Speech, Language, and Hearing Research, 51(5), 1183–1202. https://doi.org/10.1044/1092-4388(2008/07-0119)

Giglio, L., Ostarek, M., Weber, K., & Hagoort, P. (2021). Commonalities and Asymmetries in the Neurobiological Infrastructure for Language Production and Comprehension. Cerebral Cortex, 00, 1–14. https://doi.org/10.1093/CERCOR/BHAB287

Gramfort, A., Papadopoulo, T., Olivi, E., & Clerc, M. (2010). OpenMEEG: opensource software for quasistatic bioelectromagnetics. BioMedical Engineering OnLine, 9(1), 45. https://doi.org/10.1186/1475-925X-9-45

Griffiths, B. J., Mayhew, S. D., Mullinger, K. J., Jorge, J., Charest, I., Wimber, M., & Hanslmayr, S. (2019). Alpha/beta power decreases track the fidelity of stimulus-specific information. ELife, 8. https://doi.org/10.7554/eLife.49562

Guenther, F. H. (2016). Neural Control of Speech. The MIT Press.

Guenther, F. H., & Hickok, G. (2015). Neural Models of Motor Speech Control. In G. Hickok & S. L. Small (Eds.), Neurobiology of Language (pp. 725–740). Elsevier Inc. https://doi.org/10.1016/B978-0-12-407794-2.00058-4

Hanslmayr, S., Staudigl, T., & Fellner, M.-C. (2012). Oscillatory power decreases and long-term memory: the information via desynchronization hypothesis. Frontiers in Human Neuroscience, 6. https://doi.org/10.3389/fnhum.2012.00074

Hendriks, P., & Koster, C. (2010). Production/comprehension asymmetries in language acquisition. Lingua, 120(8), 1887–1897. https://doi.org/10.1016/j.lingua.2010.02.002

Hertrich, I., Dietrich, S., & Ackermann, H. (2016). The role of the supplementary motor area for speech and language processing. Neuroscience & Biobehavioral Reviews, 68, 602–610. https://doi.org/10.1016/j.neubiorev.2016.06.030

Hickok, G. (2012). Computational neuroanatomy of speech production. Nature Reviews Neuroscience, 13(2), 135–145. https://doi.org/10.1038/nrn3158

Hickok, G., Houde, J., & Rong, F. (2011). Sensorimotor Integration in Speech Processing: Computational Basis and Neural Organization. Neuron, 69(3), 407–422. https://doi.org/10.1016/j.neuron.2011.01.019

Hodapp, A., & Rabovsky, M. (2021). The N400 ERP component reflects an error-based implicit learning signal during language comprehension. European Journal of Neuroscience. https://doi.org/10.1111/ejn.15462

Huettig, F. (2015). Four central questions about prediction in language processing. Brain Research, 1626, 118–135. https://doi.org/10.1016/J.BRAINRES.2015.02.014

Jakuszeit, M., Kotz, S. A., & Hasting, A. S. (2013). Generating predictions: Lesion evidence on the role of left inferior frontal cortex in rapid syntactic analysis. Cortex, 49(10), 2861–2874. https://doi.org/10.1016/j.cortex.2013.05.014

Jenson, D., Reilly, K. J., Harkrider, A. W., Thornton, D., & Saltuklaroglu, T. (2018). Trait related sensorimotor deficits in people who stutter: An EEG investigation of µ rhythm dynamics during spontaneous fluency. NeuroImage: Clinical, 19, 690–702. https://doi.org/10.1016/J.NICL.2018.05.026

Jones, M., Gebski, V., Onslow, M., & Packman, A. (2002). Statistical Power in Stuttering Research. Journal of Speech, Language, and Hearing Research, 45(2), 243–255. https://doi.org/10.1044/1092-4388(2002/019)

Joos, K., De Ridder, D., Boey, R. A., & Vanneste, S. (2014). Functional connectivity changes in adults with developmental stuttering: a preliminary study using quantitative electro-encephalography. Frontiers in Human Neuroscience, 0(OCT), 783. https://doi.org/10.3389/FNHUM.2014.00783

Keenan, J. S. (1968). The Nature of Receptive and Expressive Impairments in Aphasia. Journal of Speech and Hearing Disorders, 33(1), 20–25. https://doi.org/10.1044/jshd.3301.20

Kronfeld-Duenias, V., Amir, O., Ezrati-Vinacour, R., Civier, O., & Ben-Shachar, M. (2016a). The frontal aslant tract underlies speech fluency in persistent developmental stuttering. Brain Structure and Function, 221(1), 365–381. https://doi.org/10.1007/s00429-014-0912-8

Kronfeld-Duenias, V., Amir, O., Ezrati-Vinacour, R., Civier, O., & Ben-Shachar, M. (2016b). Dorsal and ventral language pathways in persistent developmental stuttering. Cortex, 81, 79–92. https://doi.org/10.1016/j.cortex.2016.04.001

Kronfeld-Duenias, V., Civier, O., Amir, O., Ezrati-Vinacour, R., & Ben-Shachar, M. (2018). White matter pathways in persistent developmental stuttering: Lessons from tractography. Journal of Fluency Disorders, 55, 68–83. https://doi.org/10.1016/J.JFLUDIS.2017.09.002

Kuperberg, G. R., Brothers, T., & Wlotko, E. W. (2020). A Tale of Two Positivities and the N400: Distinct Neural Signatures Are Evoked by Confirmed and Violated Predictions at Different Levels of Representation. Journal of Cognitive Neuroscience, 32(1), 12–35. https://doi.org/10.1162/jocn_a_01465

Kuperberg, G. R., & Jaeger, T. F. (2016). What do we mean by prediction in language comprehension? Language, Cognition and Neuroscience, 31(1), 32–59. https://doi.org/10.1080/23273798.2015.1102299

Kutas, M., & Federmeier, K. D. (2011). Thirty Years and Counting: Finding Meaning in the N400 Component of the Event-Related Brain Potential (ERP). Annual Review of Psychology, 62(1), 621–647. https://doi.org/10.1146/annurev.psych.093008.131123

Lau, E. F., Phillips, C., & Poeppel, D. (2008). A cortical network for semantics: (de)constructing the N400. Nature Reviews Neuroscience, 9(12), 920–933. https://doi.org/10.1038/nrn2532

Lima, C. F., Krishnan, S., & Scott, S. K. (2016). Roles of Supplementary Motor Areas in Auditory Processing and Auditory Imagery. In Trends in Neurosciences (Vol. 39, Issue 8, pp. 527–542). Elsevier Ltd. https://doi.org/10.1016/j.tins.2016.06.003

Lukic, S., Thompson, C. K., Barbieri, E., Chiappetta, B., Bonakdarpour, B., Kiran, S., Rapp, B., Parrish, T. B., & Caplan, D. (2020). Common and Distinct Neural Substrates of Sentence Production and Comprehension. NeuroImage, 117374. https://doi.org/10.1016/j.neuroimage.2020.117374

Maris, E., & Oostenveld, R. (2007). Nonparametric statistical testing of EEG-and MEG-data. Journal of Neuroscience Methods, 164(1), 177–190. https://doi.org/10.1016/j.jneumeth.2007.03.024

Max, L., Guenther, F. H., Gracco, V. L., Ghosh, S. S., & Wallace, M. E. (2004). Unstable or Insufficiently Activated Internal Models and Feedback-Biased Motor Control as Sources of Dysfluency: A Theoretical Model of Stuttering. Contemporary Issues in Communication Science and Disorders, 31(Spring), 105–122. https://doi.org/10.1044/cicsd_31_S_105

Maxfield, N. D. (2020). Inhibitory Control of Lexical Selection in Adults who Stutter. Journal of Fluency Disorders, 105780. https://doi.org/10.1016/j.jfludis.2020.105780

McQueen, J., & Meyer, A. S. (2019). Key Issues and Future Directions: Toward a Comprehensive Cognitive Architecture for Language Use. In P. Hagoort (Ed.), Human Language: From Genes and Brains to Behavior (pp. 85–96). The MIT Press.

Mersov, A.-M., Jobst, C., Cheyne, D. O., & De Nil, L. (2016). Sensorimotor Oscillations Prior to Speech Onset Reflect Altered Motor Networks in Adults Who Stutter. Frontiers in Human Neuroscience, 10, 443. https://doi.org/10.3389/fnhum.2016.00443

Mock, J. R., Foundas, A. L., & Golob, E. J. (2016). Cortical activity during cued picture naming predicts individual differences in stuttering frequency. Clinical Neurophysiology, 127(9), 3093–3101. https://doi.org/10.1016/J.CLINPH.2016.06.005

Molinaro, N., & Monsalve, I. F. (2018). Perceptual facilitation of word recognition through motor activation during sentence comprehension. Cortex, 108, 144–159. https://doi.org/10.1016/j.cortex.2018.07.001

Molinaro, N., Monsalve, I. F., & Lizarazu, M. (2016). Is there a common oscillatory brain mechanism for producing and predicting language? Language, Cognition and Neuroscience, 31(1), 145–158. https://doi.org/10.1080/23273798.2015.1077978

Murase, S., Kawashima, T., Satake, H., & Era, S. (2016). An event-related potential investigation of sentence processing in adults who stutter. Neuroscience Research, 106, 29–37. https://doi.org/10.1016/j.neures.2015.10.004

Neef, N. E., Anwander, A., Bütfering, C., Schmidt-Samoa, C., Friederici, A. D., Paulus, W., & Sommer, M. (2018). Structural connectivity of right frontal hyperactive areas scales with stuttering severity. Brain, 141(1), 191–204. https://doi.org/10.1093/brain/awx316

Neef, N. E., Anwander, A., & Friederici, A. D. (2015). The Neurobiological Grounding of Persistent Stuttering: from Structure to Function. In Current Neurology and Neuroscience Reports (Vol. 15, Issue 9, pp. 1–11). Current Medicine Group LLC 1. https://doi.org/10.1007/s11910-015-0579-4

Neef, N. E., Bütfering, C., Anwander, A., Friederici, A. D., Paulus, W., & Sommer, M. (2016). Left posterior-dorsal area 44 couples with parietal areas to promote speech fluency, while right area 44 activity promotes the stopping of motor responses. NeuroImage, 142, 628–644. https://doi.org/10.1016/J.NEUROIMAGE.2016.08.030

Neef, N. E., Primaßin, A., Gudenberg, A. W. von, Dechent, P., Riedel, C., Paulus, W., & Sommer, M. (2021). Two cortical representations of voice control are differentially involved in speech fluency. Brain Communications, 3(2). https://doi.org/10.1093/braincomms/fcaa232

Nicenboim, B., Vasishth, S., & Rösler, F. (2020). Are words pre-activated probabilistically during sentence comprehension? Evidence from new data and a bayesian random-effects meta-analysis using publicly available data. Neuropsychologia, 142, 107427. https://doi.org/10.1016/j.neuropsychologia.2020.107427

Nieuwland, M. S., Barr, D. J., Bartolozzi, F., Busch-Moreno, S., Darley, E., Donaldson, D. I., Ferguson, H. J., Fu, X., Heyselaar, E., Huettig, F., Matthew Husband, E., Ito, A., Kazanina, N., Kogan, V., Kohút, Z., Kulakova, E., Mézière, D., Politzer-Ahles, S., Rousselet, G., … Von Grebmer Zu Wolfsthurn, S. (2020). Dissociable effects of prediction and integration during language comprehension: evidence from a large-scale study using brain potentials. Philosophical Transactions of the Royal Society B: Biological Sciences, 375(1791), 20180522. https://doi.org/10.1098/rstb.2018.0522

Oldfield, R. C. (1971). The assessment and analysis of handedness: The Edinburgh inventory. Neuropsychologia, 9(1), 97–113. https://doi.org/10.1016/0028-3932(71)90067-4

Pascual-Marqui, R. D., Michel, C. M., & Lehmann, D. (1994). Low resolution electromagnetic tomography: a new method for localizing electrical activity in the brain. International Journal of Psychophysiology, 18(1), 49–65. https://doi.org/10.1016/0167-8760(84)90014-X

Piai, V., & Zheng, X. (2019). Speaking waves: Neuronal oscillations in language production. Psychology of Learning and Motivation, 71, 265–302. https://doi.org/10.1016/bs.plm.2019.07.002

Pickering, M. J., & Gambi, C. (2018). Predicting while comprehending language: A theory and review. Psychological Bulletin, 144(10), 1002–1044. https://doi.org/10.1037/bul0000158

Pickering, M. J., & Garrod, S. (2013). An integrated theory of language production and comprehension. Behavioral and Brain Sciences, 36(4), 329–347. https://doi.org/10.1017/S0140525X12001495

Prystauka, Y., & Lewis, A. G. (2019). The power of neural oscillations to inform sentence comprehension: A linguistic perspective. Language and Linguistics Compass, 13(9). https://doi.org/10.1111/lnc3.12347

Riley, G. D. (2009). The stuttering severity instrument for adults and children (SSI-4). 4th Edition. PRO-ED Inc.

Rommers, J., Dickson, D. S., Norton, J. J. S., Wlotko, E. W., & Federmeier, K. D. (2017). Alpha and theta band dynamics related to sentential constraint and word expectancy. Language, Cognition and Neuroscience, 32(5), 576–589. https://doi.org/10.1080/23273798.2016.1183799

Saltuklaroglu, T., Bowers, A., Harkrider, A. W., Casenhiser, D., Reilly, K. J., Jenson, D. E., & Thornton, D. (2018). EEG mu rhythms: Rich sources of sensorimotor information in speech processing. Brain and Language, 187, 41–61. https://doi.org/10.1016/J.BANDL.2018.09.005

Shalom, D. Ben, & Poeppel, D. (2008). Functional anatomic models of language: assembling the pieces. The Neuroscientist : A Review Journal Bringing Neurobiology, Neurology and Psychiatry, 14(1), 119–127. https://doi.org/10.1177/1073858407305726

Silbert, L. J., Honey, C. J., Simony, E., Poeppel, D., & Hasson, U. (2014). Coupled neural systems underlie the production and comprehension of naturalistic narrative speech. Proceedings of the National Academy of Sciences, 111(43), E4687–E4696. https://doi.org/10.1073/pnas.1323812111

Siman-Tov, T., Granot, R. Y., Shany, O., Singer, N., Hendler, T., & Gordon, C. R. (2019). Is there a prediction network? Meta-analytic evidence for a cortical-subcortical network likely subserving prediction. Neuroscience & Biobehavioral Reviews, 105, 262–275. https://doi.org/10.1016/j.neubiorev.2019.08.012

Smith, A., & Weber, C. (2017). How Stuttering Develops: The Multifactorial Dynamic Pathways Theory. Journal of Speech, Language, and Hearing Research, 60(9), 2483–2505. https://doi.org/10.1044/2017_JSLHR-S-16-0343

Tadel, F., Baillet, S., Mosher, J. C., Pantazis, D., & Leahy, R. M. (2011). Brainstorm: a user-friendly application for MEG/EEG analysis. Computational Intelligence and Neuroscience, 2011, 879716. https://doi.org/10.1155/2011/879716

Tourville, J. A., & Guenther, F. H. (2011). The DIVA model: A neural theory of speech acquisition and production. Language and Cognitive Processes, 26(7), 952–981. https://doi.org/10.1080/01690960903498424

Tremblay, P., & Dick, A. S. (2016). Broca and Wernicke are dead, or moving past the classic model of language neurobiology. Brain and Language, 162, 60–71. https://doi.org/10.1016/j.bandl.2016.08.004

Urbach, T. P., DeLong, K. A., Chan, W.-H., & Kutas, M. (2020). An exploratory data analysis of word form prediction during word-by-word reading. Proceedings of the National Academy of Sciences, 117(34), 201922028. https://doi.org/10.1073/pnas.1922028117

van Casteren, M., & Davis, M. H. (2006). Mix, a program for pseudorandomization. Behavior Research Methods, 38(4), 584–589. https://doi.org/10.3758/BF03193889

Walenski, M., Europa, E., Caplan, D., & Thompson, C. K. (2019). Neural networks for sentence comprehension and production: An ALE-based meta-analysis of neuroimaging studies. Human Brain Mapping, 40(8), 2275–2304. https://doi.org/10.1002/hbm.24523

Wang, L., Hagoort, P., & Jensen, O. (2018). Language Prediction Is Reflected by Coupling between Frontal Gamma and Posterior Alpha Oscillations. Journal of Cognitive Neuroscience, 30(3), 432–447. https://doi.org/10.1162/jocn_a_01190

Weber-Fox, C. (2001). Neural Systems for Sentence Processing in Stuttering. Journal of Speech, Language, and Hearing Research, 44(4), 814–825. https://doi.org/10.1044/1092-4388(2001/064)

Weber-Fox, C., & Hampton, A. (2008). Stuttering and Natural Speech Processing of Semantic and Syntactic Constraints on Verbs. Journal of Speech, Language, and Hearing Research. 1 In the literature, PWS is used to refer to both adults and children who stutter (AWS and CWS, respectively) indistinctively. The appropriate acronyms are used accordingly throughout the manuscript. https://doi.org/10.1044/1092-4388(2008/07-0164)

