## Supplementary Material (PDF) for "When inefficient speech-motor control affects speech comprehension: atypical electrophysiological correlates of language prediction in stuttering"

#### A) SUPPLEMENTARY TABLES AND FIGURES

| | LISTS | | | | |
|---|---|---|---|---|---|
| | **Mean (sd) List A** | **Mean (sd) List B** | ***t*-value** | **df** | ***p*-value** |
| **Lexical frequency (log-scaled)** | 3.776 (1.252) | 3.849 (1.305) | -0.46 | 235.56 | 0.648 |
| **No. phonemes (word)** | 6.063 (1.701) | 6.281 (1.631) | -1.05 | 253.55 | 0.295 |
| **No. syllables (word)** | 2.578 (0.728) | 2.672 (0.641) | -1.09 | 250.06 | 0.275 |
| **No. syllables (sentence frame)** | 10.680 (2.012) | 10.672 (2.248) | 0.03 | 251.07 | 0.977 |
| **Audio length (sec) (sentence frame)** | 2.388 (0.397) | 2.381 (0.414) | 0.15 | 253.54 | 0.885 |
| **No. syllables/sec (sentence frame)** | 4.482 (0.485) | 4.480 (0.523) | 0.05 | 252.52 | 0.964 |
| **Cloze probability overall** | 0.456 (0.421) | 0.468 (0.421) | -0.23 | 245 | 0.822 |
| **Cloze probability HC** | 0.868 (0.094) | 0.878 (0.09) | -0.62 | 125.84 | 0.535 |
| **Cloze probability LC** | 0.045 (0.068) | 0.058 (0.086) | -1 | 119.69 | 0.319 |
| | **CONDITIONS** | | | | |
| | **Mean (sd) HC** | **Mean (sd) LC** | ***t*-value** | **df** | ***p*-value** |
| **No. syllables (sentence frame)** | 10.750 (2.074) | 10.602 (2.182) | 0.56 | 253.34 | 0.577 |
| **Audio length (sec) (sentence frame)** | 2.413 (0.407) | 2.356 (0.402) | 1.12 | 253.97 | 0.263 |
| **No. syllables/sec** | 4.459 (0.441) | 4.504 (0.56) | -0.71 | 240.87 | 0.476 |
| **Cloze probability overall** | 0.873 (0.092) | 0.052 (0.077) | 77.47 | 246.66 | <0.001 |
| **Cloze probability List A** | 0.868 (0.094) | 0.045 (0.068) | 56.96 | 114.71 | <0.001 |
| **Cloze probability List B** | 0.878 (0.09) | 0.058 (0.086) | 52.69 | 125.62 | <0.001 |

**Table S1: Variables controlled across lists and conditions (Welch's t-tests).** Means and standard deviations (in parenthesis) are reported. HC: high constraint, LC: low constraint.

| AWS participant ID | SSI score | Percentile | Severity |
|---|---|---|---|
| 1 | 24 | 24-40 | Mild |
| 2 | 13 | 5-11 | Very mild |
| 3 | 12 | 1-4 | Very mild |
| 4 | 33 | 78-88 | Severe |
| 5 | 10 | 1-4 | Very mild |
| 6 | 24 | 24-40 | Mild |
| 7 | 16 | 5-11 | Very mild |
| 8 | 11 | 1-4 | Very mild |
| 9 | 36 | 89-95 | Severe |
| 10 | 32 | 78-88 | Severe |
| 11 | 22 | 24-40 | Mild |
| 12 | 16 | 5-11 | Very mild |
| 13 | 21 | 24-40 | Mild |
| 14 | 18 | 12-33 | Mild |

**Table S2: SSI-4 scores for each participant.**

<table>
<tr><th>Custom ROI number</th><th>Custom atlas</th><th>Desikan-Killiany atlas</th></tr>
<tr><td rowspan="2">1</td><td rowspan="2">Posterior frontal cortex</td><td>Caudal middle frontal</td></tr>
<tr><td rowspan="2">Superior frontal</td></tr>
<tr><td rowspan="2">2</td><td rowspan="2">Anterior frontal cortex</td></tr>
<tr><td>Rostral middle frontal</td></tr>
<tr><td rowspan="3">3</td><td rowspan="3">Inferior frontal cortex</td><td>Pars orbitalis</td></tr>
<tr><td>Pars opercularis</td></tr>
<tr><td>Pars triangularis</td></tr>
<tr><td>4</td><td>Precentral gyrus</td><td>Precentral</td></tr>
<tr><td>5</td><td>Postcentral gyrus</td><td>Postcentral</td></tr>
<tr><td>6</td><td>Insula</td><td>Insula</td></tr>
<tr><td>7</td><td>Superior parietal cortex</td><td>Superior parietal</td></tr>
<tr><td rowspan="2">8</td><td rowspan="2">Inferior parietal cortex</td><td>Supramarginal</td></tr>
<tr><td>Inferior parietal</td></tr>
<tr><td rowspan="6">9</td><td rowspan="6">Temporal cortex</td><td>Transverse temporal</td></tr>
<tr><td>Superior temporal</td></tr>
<tr><td>Middle Temporal</td></tr>
<tr><td>Inferior temporal</td></tr>
<tr><td>Temporal pole</td></tr>
<tr><td>Banks of Superior temporal sulcus</td></tr>
</table>

**Table S3: Custom atlas with 9 regions of interest (ROIs) defined starting from the Desikan-Killiany parcellation (Desikan et al., 2006).** Each ROI has been defined for the left and the right hemisphere. Some regions were merged into one single ROI (e.g., pars orbitalis, pars triangularis and pars opercularis merged into the inferior frontal gyrus). The merge was performed in order to reduce the number of ROIs, also considering the spatial inaccuracy of the EEG technique that would not differentiate in a significant way the signal stemming from very close cortical sources (e.g., pars triangularis and pars opercularis). In one case, a unitary region in the Desikan-Killiany parcellation (i.e., superior frontal cortex ROI) was subdivided in 6 sub-regions, by using a dedicated function in Brainstorm. Subsequently, the caudal sub-regions were merged with the caudal middle frontal region, defining the posterior frontal cortex, while the rostral sub-regions were merged with the rostral middle frontal region, defining the anterior frontal cortex. The rationale behind this decision was the necessity to identify pre-motor associative regions, that may include crucial areas involved in stuttering (e.g., the SMA-complex; see Busan, 2020; Chang & Guenther, 2020; Civier, Bullock, Max, & Guenther, 2013), with respect to pre-frontal regions, usually more associated to executive functions.

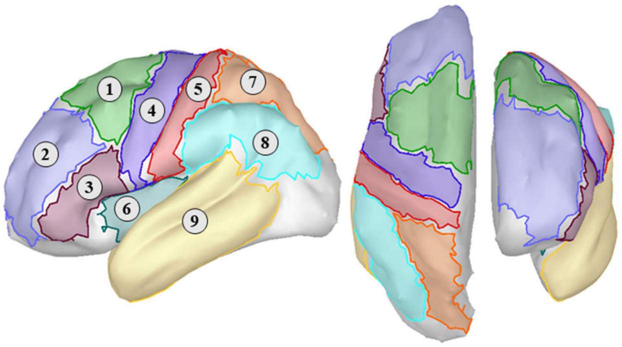


**Figure S1: ROIs projected on an inflated cortical model.** For the sake of simplicity, the left hemisphere only is visualized. Lateral (left), dorsal (middle), and rostral (right) views of the cortical model are displayed. 1: posterior frontal cortex; 2: anterior frontal cortex; 3: inferior frontal cortex; 4: precentral gyrus; 5: postcentral gyrus; 6: insula; 7: superior parietal cortex; 8: inferior parietal cortex; 9: temporal cortex.

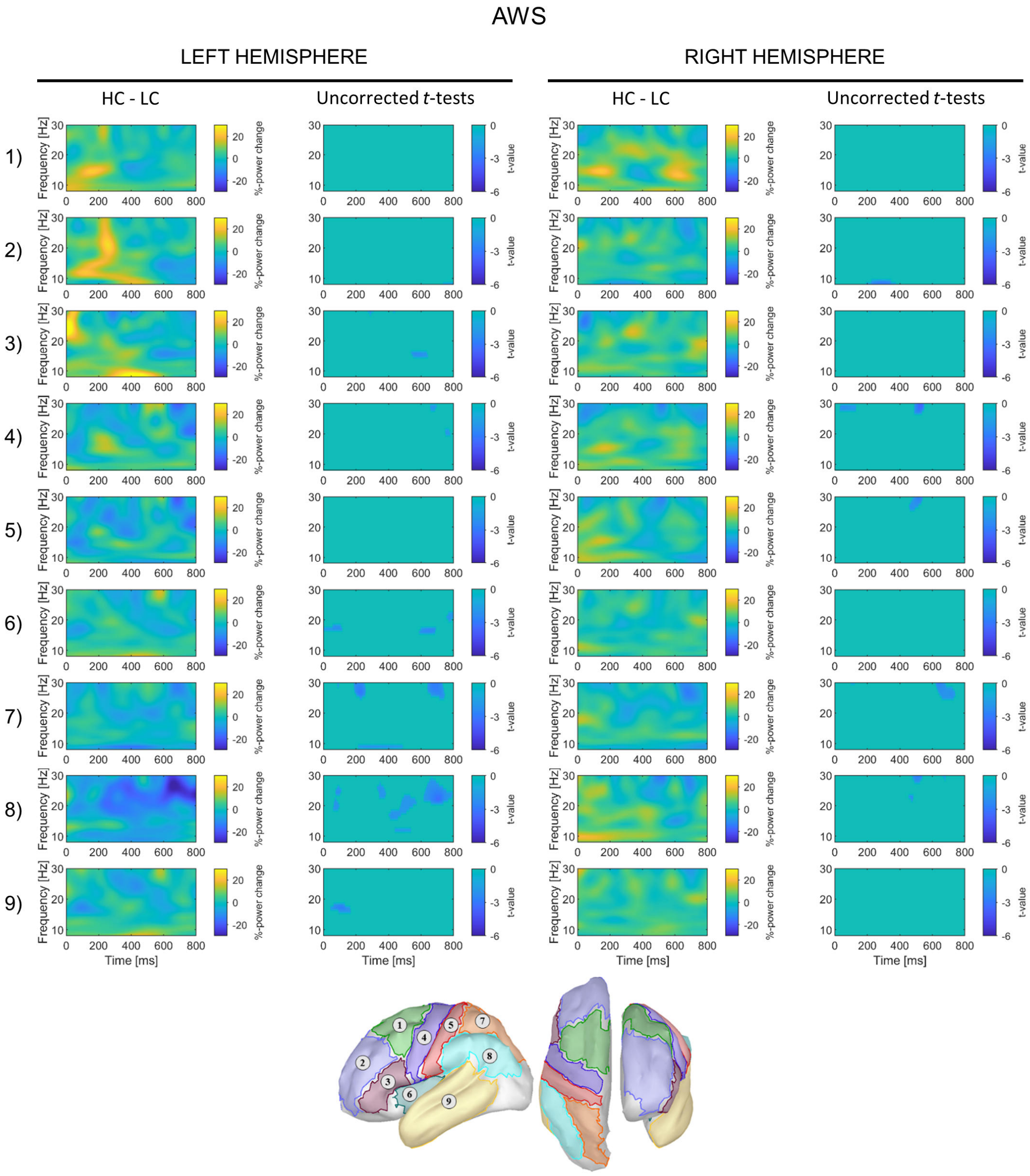


**Figure S2 | Time-frequency results at the ROI level, comprehension task, AWS**

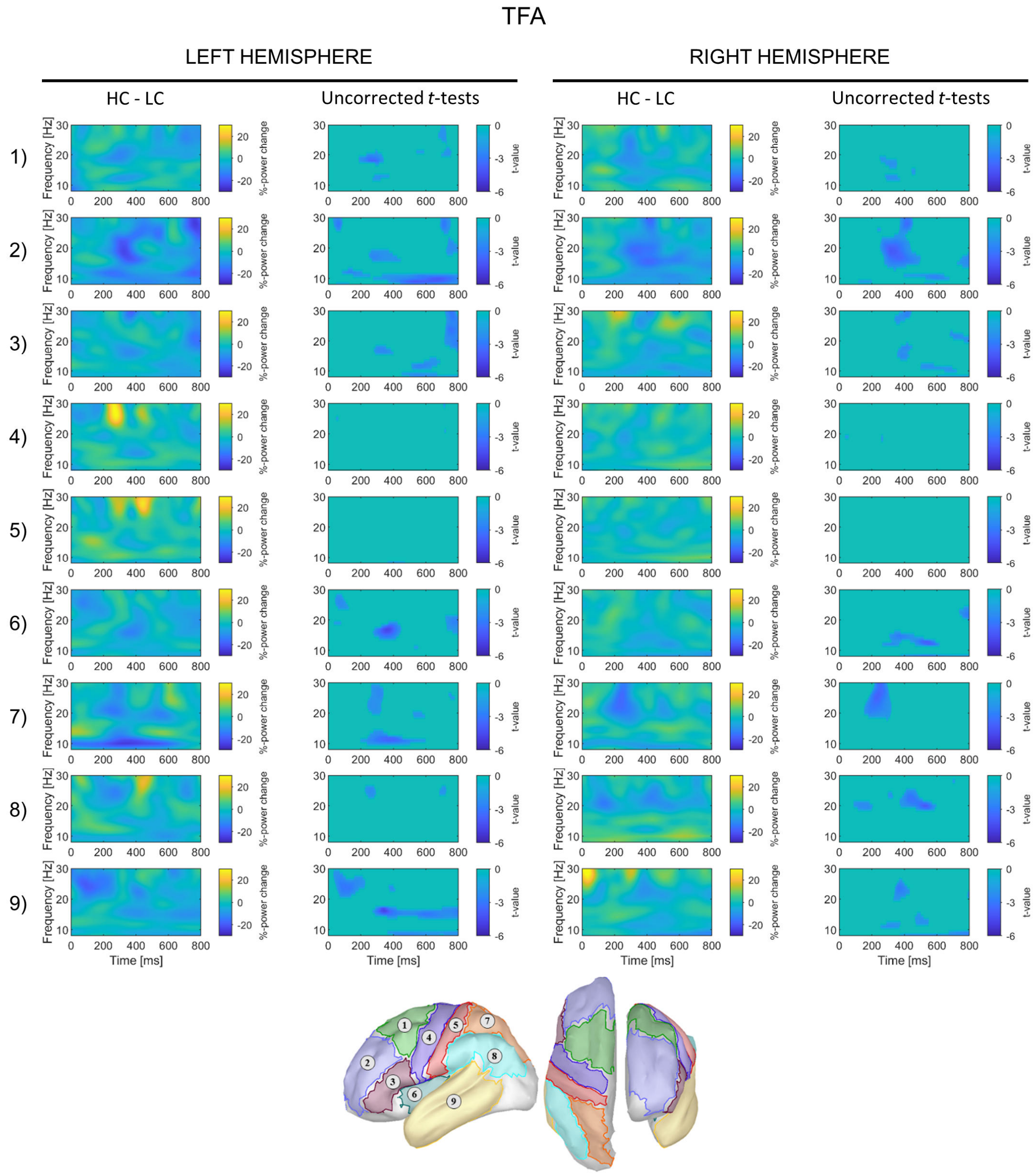


**Figure S3 | Time-frequency results at the ROI level, comprehension task, TFA**

### B) PRODUCTION TASK

#### Introduction

Alpha–beta (especially beta) activity is prominently associated to sensorimotor processes and action control. Typically, beta power in sensorimotor regions decreases during movement planning and execution, with an increase (rebound) right after movement completion (Kilavik et al., 2013). Beta power decrease is strictly related to motor readiness, and it correlates with the likelihood of initiating a voluntary action (Jenkinson & Brown, 2011). Specifically in the speech-motor control domain, pre-speech onset motor-specific beta power decrease is thought to reflect the generation of top-down predictions via forward models, which send sensory predictions to the auditory cortex for self-monitoring (Saltuklaroglu et al., 2018). Outside the strictly motor aspect, beta power decrease has been found to be involved also in memory aspects of language production. In context-induced picture naming tasks, for instance, participants listen to sentences and can plan the production of a word (the linguistic label of a picture) before the picture is displayed when the context is highly constraining. Alpha–beta power decreases are consistently found before picture onset in constraining conditions, reflecting the engagement of production processes and the activation of linguistic (and possibly motor) information for word planning and execution. Additionally, participants are consistently faster in their responses in high constraining conditions (Gastaldon et al., 2020; Piai et al., 2014, 2015, 2018, 2020).

In developmental stuttering (DS), these frequencies have been found to be atypically modulated, reflecting disrupted sensorimotor mechanisms. For instance, Jenson et al. (2018) investigated syllable and word repetition in DS, showing a reduced power decrease in the beta band in the sensorimotor cortices, signaling weaker forward modeling or reduced confidence in sensory predictions. In a sentence repetition task, Mersov et al. (2016) found the opposite modulation, namely stronger beta power decrease in left premotor areas, reflecting reduced coordination in the motor network when planning speech production. Compatibly, Mock et al. (2016) found that, in a cued word production task, the degree of beta decrease positively correlated with stuttering rate. Overall, studies suggest that activity in the beta band is differently modulated in PWS (see also Joos, Ridder, Boey, & Vanneste, 2014), reflecting difficulties in setting up and coordinating activity in neural structures involved in speech planning and execution.

In this part of the study, we capitalize on these findings (alpha–beta power decrease in speech planning and execution, and atypical beta modulations in DS) to explore, with a context-induced picture naming task, whether anticipating word planning in AWS is reflected in atypical alpha–beta power modulations, in a paradigm that triggers production planning in a different way relative to traditional studies on speech production in stuttering (i.e., internally generated *vs.* externally induced).

#### Methods

##### *Procedure*

As in the comprehension task, in the production task participants were presented with a fixation cross (800 ms) before the auditory sentence frame was presented (while still displaying the fixation cross). After a silent gap of 800 ms, a target picture was displayed for 2 seconds. Participants were asked to name the picture as quickly and as accurately as possible (see Figure S4). The vocal responses were recorded over the same interval of the picture presentation (2 s). All responses were collected via a microphone, positioned at a fixed distance from the participant (~50 cm). Trial order was pseudo-randomized for each participant as described in the main text of the manuscript. Participants were allowed to take a break every 32 minutes. A familiarization phase was carried out before the task (8 trials, no stimuli were presented in the subsequent experiment). The task lasted approximately 20 minutes.

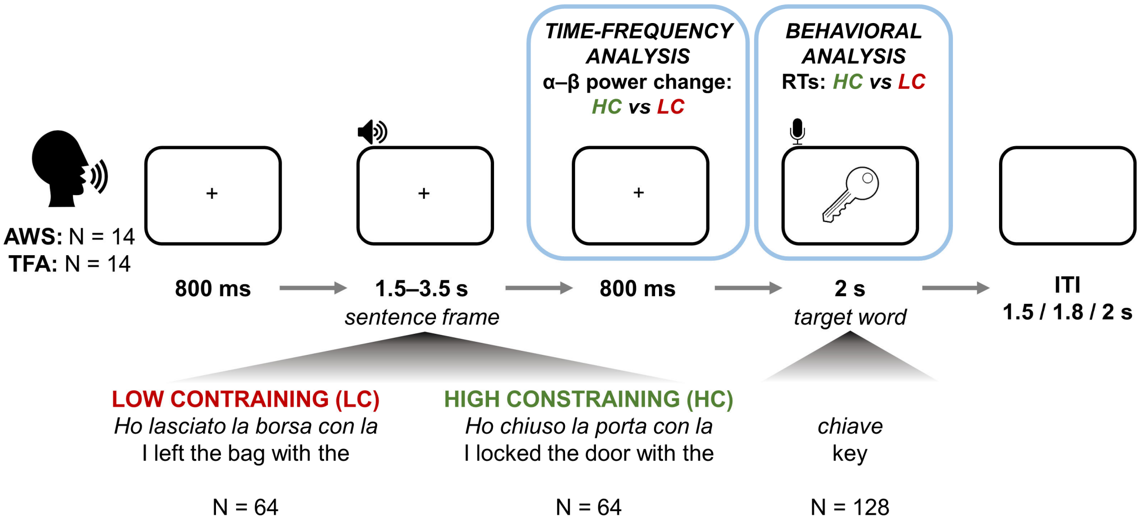


**Figure S4 | Task structure and analyses of interest (production)**

##### *Behavioral response coding and statistical analysis*

The recordings were individually inspected, and responses were manually coded as incorrect when participants: 1) failed to provide an answer within the 2-second window, 2) produced hesitation sounds (e.g., 'ehm', 'mmm') before providing the answer, 3) started producing a word but then produced another word, 4) started producing a word before recording onset, thus making response time (RT) measurement inadequate. For the AWS group, stuttering events (blocks, repetitions, prolongations) prior to a correct response were not considered as a criterion for response exclusion, since these do not automatically signal a difficulty in retrieving the linguistic labels. However, overt stuttering events in AWS were occasional and very rare during the experimental trials. Trials with incorrect responses were excluded from the EEG analyses. RTs were calculated by submitting the individual recordings to Chronset (Roux et al., 2017). In case Chronset returned some NA values, the correspondent audio waveforms were inspected manually with Audacity in order to determine the response onset. The set of correct responses was then analyzed using R (R Core Team, 2018), by means of linear mixed effects models of increasing complexity. First a model with only random effects (intercept for participant and target word) was computed; each subsequent model included a predictor or an interaction (see Table S5). A model comparison approach was adopted to select the model with the best fit: an ANOVA between models was performed, and the AIC and the p-value were taken as indices of fit. Then, a type III ANOVA of the best model was performed to obtain estimates of the effects. To explore interactions between predictors highlighted by the best model, post-hoc pairwise contrasts between estimated marginal effects (Tukey's tests) were carried out with the R package *emmeans* (Lenth, 2020).

##### *EEG pre-processing and analyses*

All pre-processing and analyses steps for the production task were the same as for the comprehension task, as described in the main text of the manuscript. The mean percentage of epochs retained for each group are the following: AWS: baseline: 88%, $\text{interval}_{HC}$: 87.5%, $\text{interval}_{LC}$: 87.2%; TFA: baseline: 86.8%, $\text{interval}_{HC}$: 89.2%, $\text{interval}_{LC}$: 87.6%.

##### *Correlations*

Correlations were performed only on the ROIs that highlighted significant clusters in the within-group comparisons. Power averages were computed as described for the comprehension task (see main text). These power changes (Δ%power-change) were correlated with ΔRTs, that is, the difference in subject-averaged response times between the HC and the LC conditions, highlighting the facilitation effect of predictable targets. For the AWS group, Δ%power-changes were also correlated with SSI-4 scores.

#### Results

##### *Behavioral analysis*

Accuracy was very high for both groups, but higher in TFA (AWS: 95.87%, TFA: 98.88%). A higher error rate was found for the AWS (N = 74) than for the TFA group (N = 20). In both groups, the error rate was higher in the LC condition (AWS: HC: 30, LC: 44; TFA: HC: 6, LC: 14). The higher number of incorrect responses in the AWS group, rather than reflecting lexical difficulties or lack of attention to the task, is more likely due to the 2-second time-window for providing a response and the exclusion of responses that were not provided within the recording time (i.e., the segment of speech recorded was not sufficient to evaluate the correctness of the response). Error rates were not further analyzed. See Tables S4 and Figure S5 for descriptive statistics and graphical representations.

| | **RTs mean [ms] (sd)** | | |
|---|---|---|---|
| | **HC** | **LC** | **HC - LC** |
| **AWS** | 668 (263.06) | 841 (227.92) | -173 |
| **TFA** | 548 (191.67) | 748 (183.44) | -200 |
| **AWS - TFA** | 120 | 93 | |

**Table S4: Descriptive statistics of RTs for word production (correct trials only).**

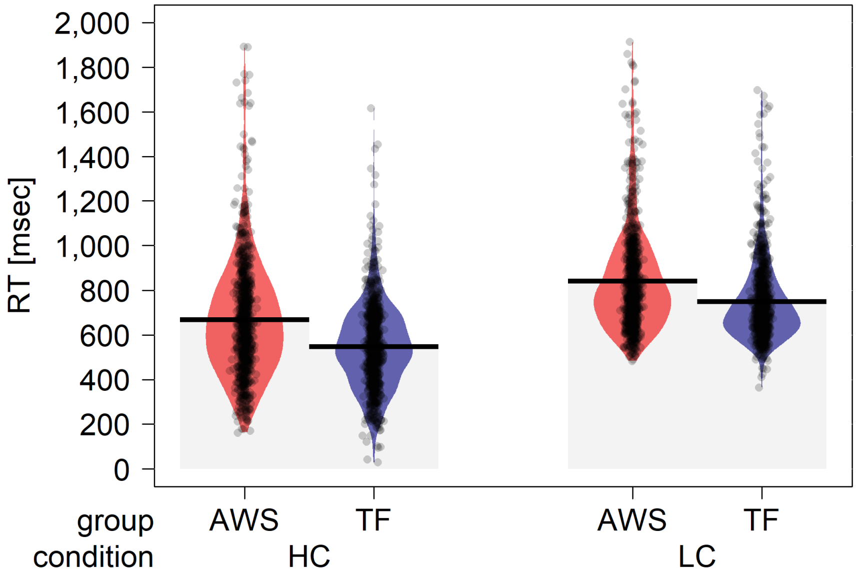


**Figure S5 | Violin plots displaying RT distributions divided by condition and group.**

In Table S5 the results of the ANOVA between models are reported, showing that the best model is *model 6*. The estimates of the fixed effects of *model 6* are shown in Table S6, and the results of the ANOVA are shown in Table S7. The model revealed main effects of repetition, group, and condition. The HC condition induces faster responses relative to the LC condition. The effect of lexical frequency emerged only in the LC condition, as revealed by the interaction: higher lexical frequency is associated to faster responses only in the LC condition. A significant interaction Group × Condition was also found. The pairwise contrasts, shown in Table S8, revealed that the effect of group is significant in the HC but not in the LC condition, highlighting faster responses for TFA relative to AWS.

| Model | Effects | AIC | $X^2$ | *p*-value |
|---|---|---|---|---|
| 0 | Random Effects (RE) | 47249 | | |
| 1 | + Repetition | 47138 | 112.65 | $< 0.001$ |
| 2 | + Group | 47135 | 5.505 | 0.019 |
| 3 | + Condition | 46261 | 875.763 | $< 0.001$ |
| 4 | + Lexical Frequency | 46260 | 2.915 | 0.088 |
| 5 | + Group × Condition | 46256 | 6.202 | 0.013 |
| 6 | + Lexical Frequency × Condition | 46250 | 7.673 | 0.006 |
| 7 | + Lexical Frequency × Condition × Group | 46252 | 2.649 | 0.266 |

**Table S5 | Model comparison.** Random Effects (RE) include random intercepts for participant and target word. At each step a predictor or an interaction is added. The best model is *model 6*.

| | Estimate | S. E. | Df | *t*-value | *p*-value | 95% CI |
|---|---|---|---|---|---|---|
| **Intercept** | 712.863 | 37.141 | 52.59 | 19.194 | < 0.001 | [638.683 – 787.064] |
| **Repetition** | -74.12 | 5.945 | 3341.484 | -12.467 | < 0.001 | [-85.776 – -62.464] |
| **Group** | -125.08 | 45.017 | 28.912 | -2.779 | 0.01 | [-216.345 – -33.836] |
| **Condition** | 222.936 | 19.758 | 3342.640 | 11.283 | < 0.001 | [184.199 – 261.672] |
| **Lexical Frequency** | -0.864 | 4.789 | 213.315 | -0.18 | 0.857 | [-10.288 – 8.57] |
| **Group × Condition** | 29.480 | 11.882 | 3339.36 | 2.481 | 0.013 | [6.185 – 52.775] |
| **Condition × Lexical Frequency** | -12.913 | 4.659 | 3341.837 | -2.772 | 0.006 | [-22.048 – -3.779] |

**Table S6 | Estimates of the fixed effects of *model 6***

| Fixed effect | *F*-value | *p*-value |
|---|---|---|
| **Repetition** | 155.431 | < 0.001 |
| **Group** | 6.113 | 0.02 |
| **Condition** | 160.048 | < 0.001 |
| **Lexical Frequency** | 3.033 | 0.084 |
| **Group × Condition** | 6.156 | 0.013 |
| **Condition × Lexical Frequency** | 7.682 | 0.006 |

**Table S7 | Type III ANOVA of model 6 (Satterthwaite's method)**

| Contrast | Estimate | SE | z-ratio | *p*-value |
|---|---|---|---|---|
| **AWS HC – TFA HC** | 125.1 | 45.02 | 2.779 | 0.028 |
| **AWS HC – AWS LC** | -173.5 | 8.47 | -20.494 | <0.0001 |
| **AWS HC – TFA LC** | -77.9 | 45.02 | -1.731 | 0.307 |
| **TFA HC – AWS LC** | -298.6 | 45.02 | -6.633 | <0.0001 |
| **TFA HC – TFA LC** | -203.0 | 8.34 | -24.356 | <0.0001 |
| **AW LC – TFA LC** | 95.6 | 45.03 | 2.123 | 0.146 |

**Table S8 | Pairwise contrasts.** Values are averaged over the level of Repetition. Degrees-of-freedom: asymptotic (p-values adjusted with Tukey method)

##### *Time-frequency analyses*

At the sensor-level, significant negative clusters have been found in the within-group contrast (AWS: $p$ = 0.004, $t$-sum = -223566, size = 92164; TFA: $p$ = 0.001, $t$-sum = -781735, size = 28418), but not in the between-group contrast. Figure S4 summarizes the results and provides cluster-corrected and uncorrected statistics.

At the ROI level, the cluster-corrected tests revealed several significant clusters when contrasting the HC and the LC conditions within each group (Table S9, Figure S5 and S6), but not when contrasting the two groups (Figure S7). For this latter case, uncorrected t-tests only are provided.

More specifically, sensor analyses resulted in diffused weaker decrease (especially in the beta range in both hemispheres) and more restricted stronger decrease in AWS (especially in the alpha range in the right hemisphere). When considering ROIs analyses, AWS showed stronger power decreases in the alpha and high beta ranges in various regions of both hemispheres, more prominently in bilateral premotor associative and prefrontal cortices, the right inferior frontal cortex, and the left somatosensory cortex. On the other hand, reduced power suppressions in the alpha and low beta range are evident in the right prefrontal cortex, right inferior parietal cortex, and the left temporal cortex.

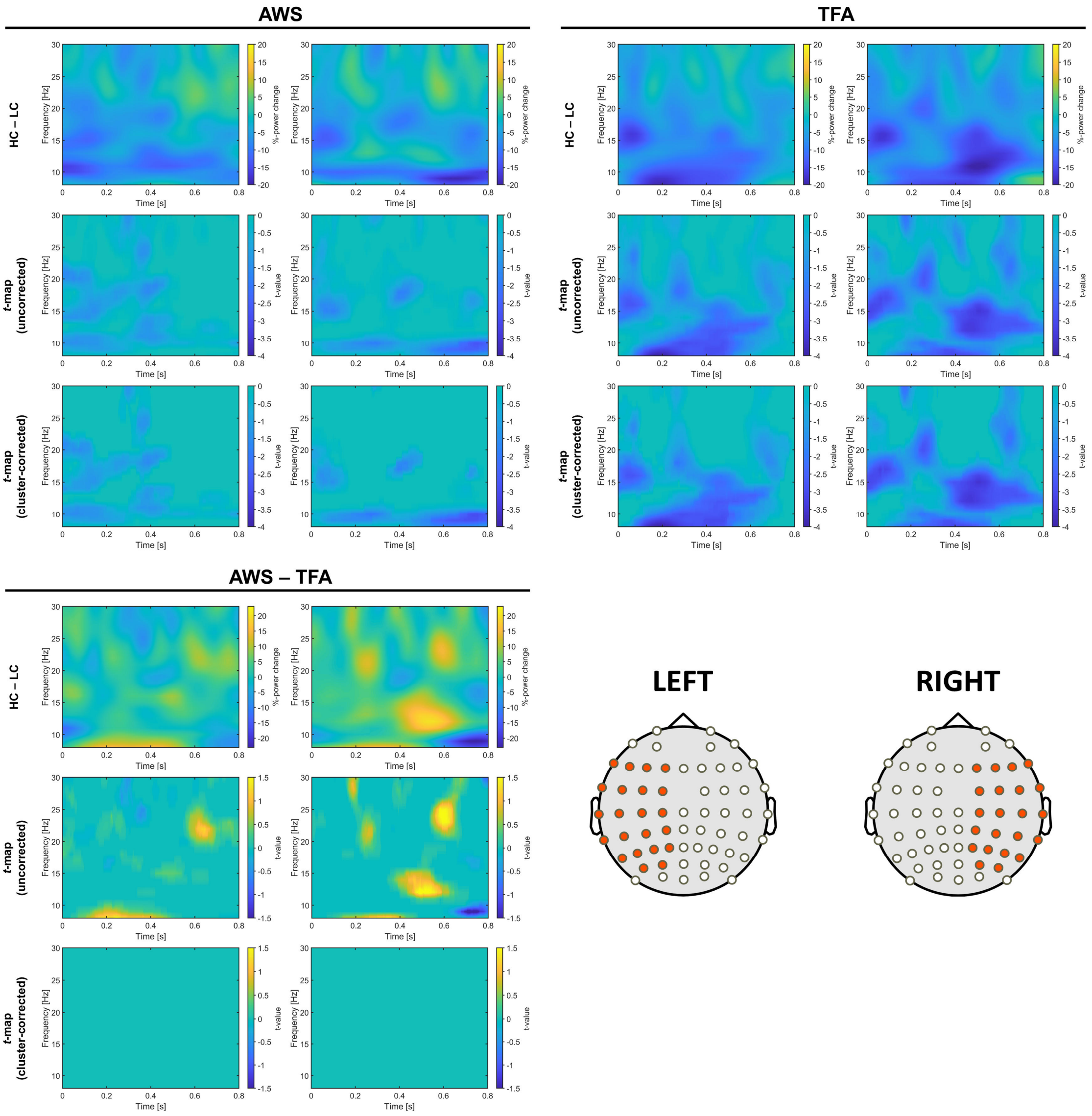


**Figure S6 | Time-frequency results at the sensor level**

| ROI | AWS | | | | | | TFA | | | | | |
|---|---|---|---|---|---|---|---|---|---|---|---|---|
| | Left | | | Right | | | Left | | | Right | | |
| | *p*-value | *t*-sum | size | *p*-value | *t*-sum | size | *p*-value | *t*-sum | size | *p*-value | *t*-sum | size |
| 1 | 0.011<br>0.028 | -20343<br>-15709 | 6142<br>5415 | 0.001 | -35328 | 12220 | | | | 0.041 | -11588 | 4761 |
| 2 | | | | | | | | | | | | |
| 3 | | | | | | | | | | | | |
| 4 | | | | 0.003 | -24310 | 9067 | | | | 0.001 | -28309 | 10000 |
| 5 | | | | 0.027 | -15936 | 5887 | | | | 0.002 | -25572 | 10263 |
| 6 | | | | | | | | | | 0.033 | -12222 | 4844 |
| 7 | | | | | | | | | | | | |
| 8 | 0.01798 | -18318 | 6785 | | | | 0.042 | -11514 | 3903 | 0.025 | -14000 | 4300 |
| 9 | | | | | | | 0.041 | -11612 | 4482 | 0.026 | -13822 | 5092 |

**Table S9 | Cluster-statistics at the ROI level.**

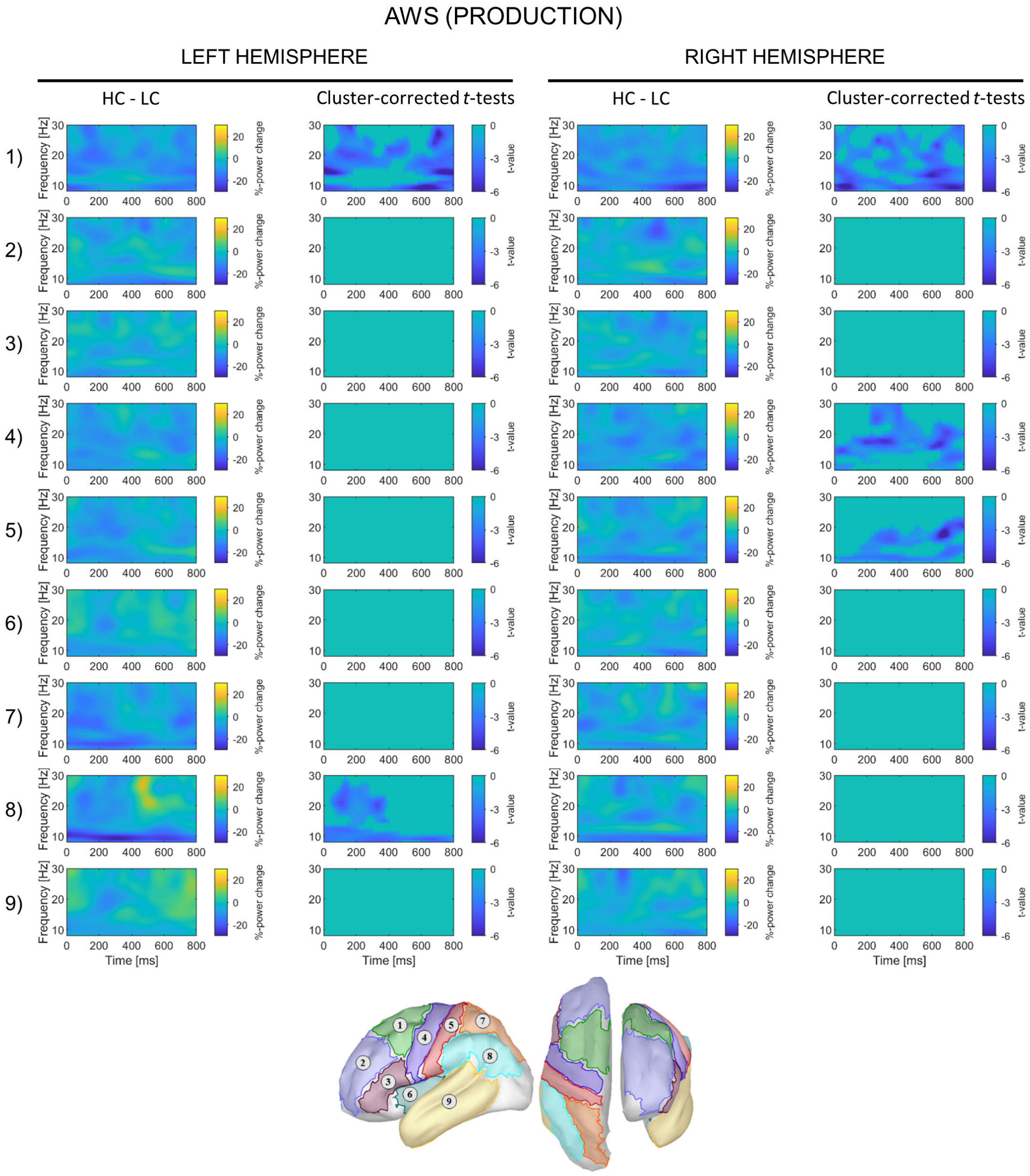


**Figure S7 | Time-frequency results at the ROI level, production task, AWS.**

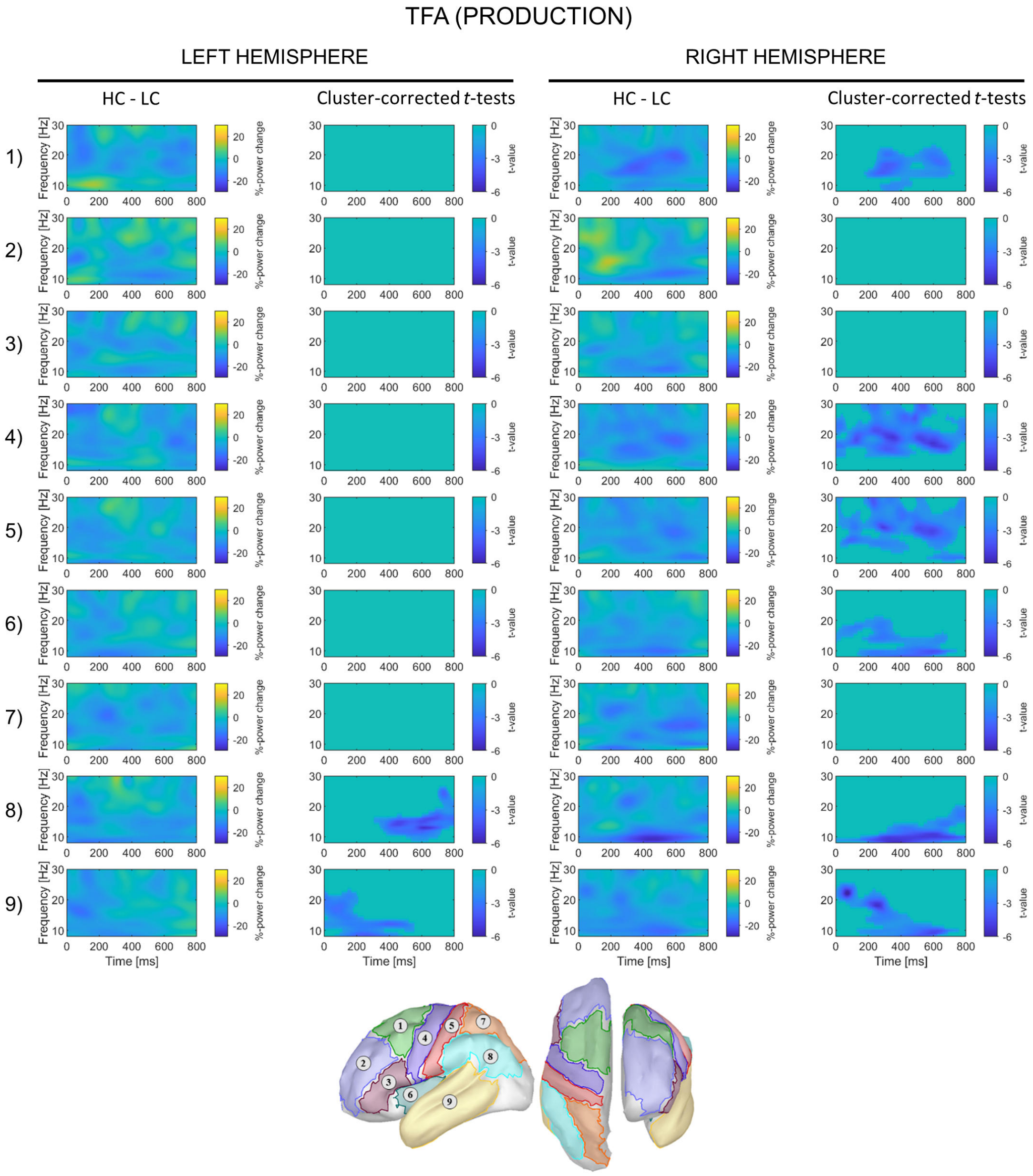


**Figure S8 | Time-frequency results at the ROI level, production task, TFA.**

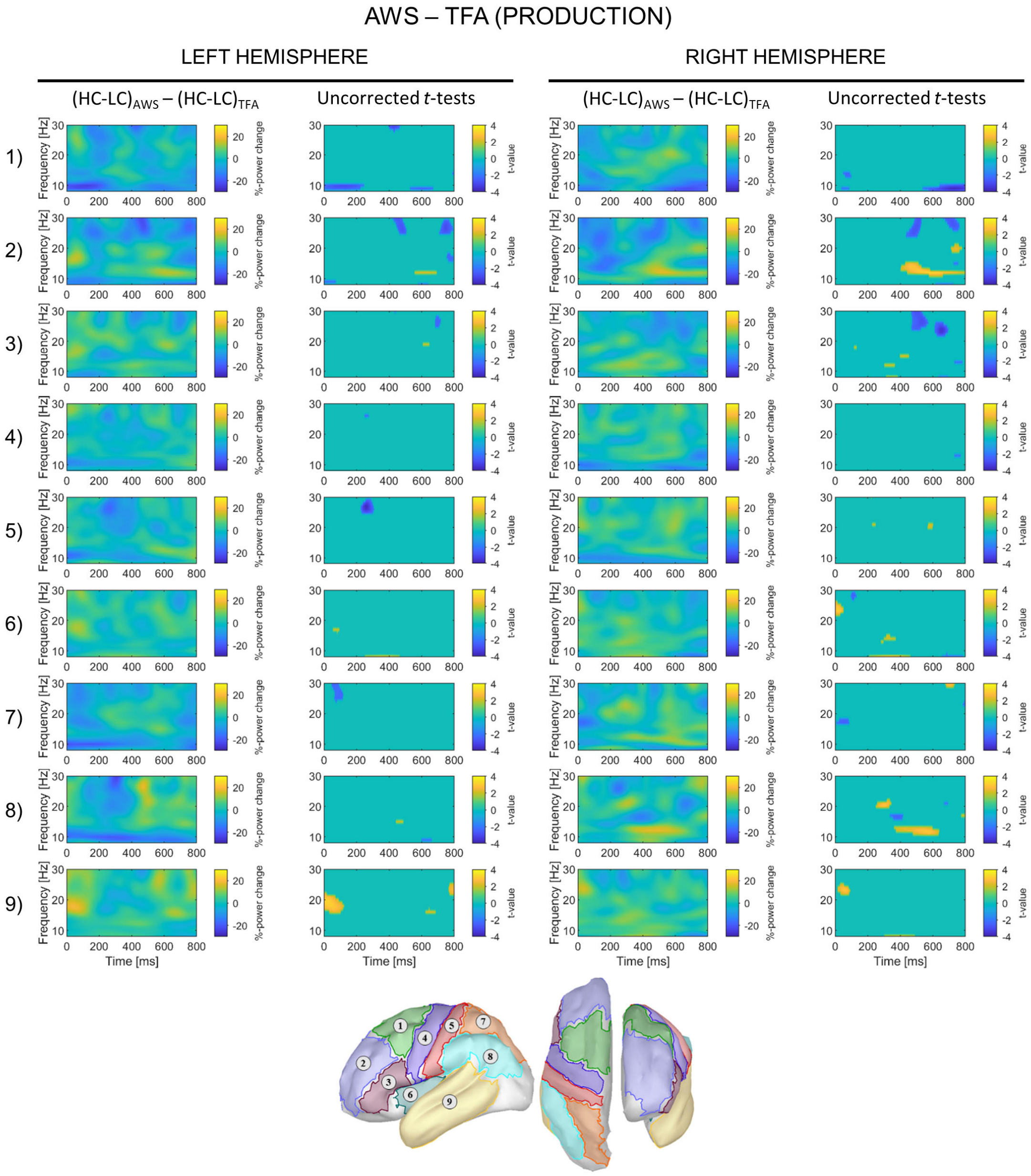


**Figure S9 | Time-frequency results at the ROI level, production task, AWS *vs.* TFA.**

##### *Correlations*

Statistically significant correlations are shown in Figure S8. In the AWS group, SSI-4 values are negatively correlated with beta1 Δ%-power change in the left posterior frontal cortex (ROI1_L) ($R$ = -0.62, $p$ = 0.019): a stronger decrease is associated to more severe stuttering. ΔRTs are positively correlated with beta2 Δ%-power change in the right motor cortex (ROI4_R) ($R$ = -0.63, $p$ = 0.016): a stronger facilitation (more negative ΔRTs) is associated to a stronger power decrease in this area.

In the TFA group, ΔRTs are positively correlated with alpha Δ%-power change in the left inferior parietal region (ROI8_L) ($R$ = 0.68, $p$ = 0.007): a stronger facilitation is associated to a stronger power decrease.

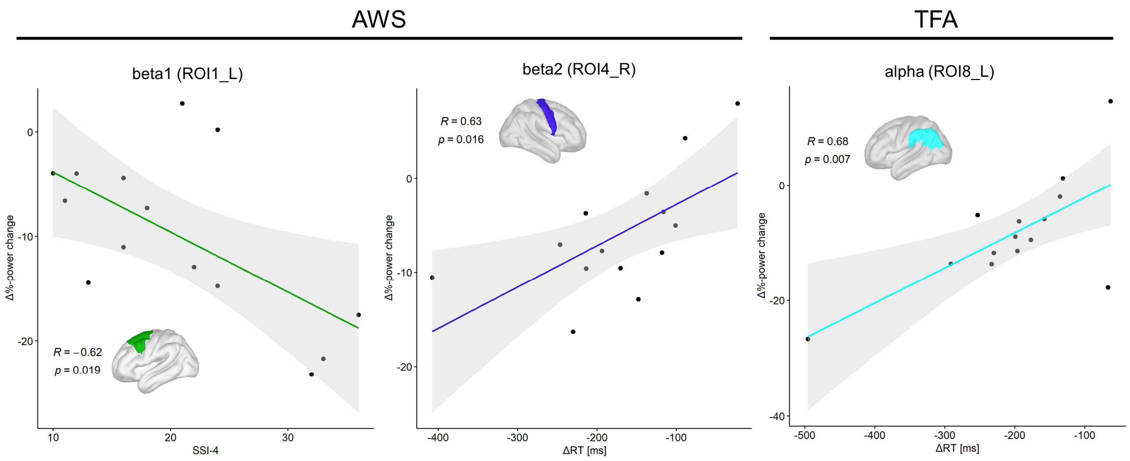


**Figure S10 | Correlations in the production task**

#### Discussion

In this part of the study, we investigated oscillatory and behavioral correlates of context-induced word production in AWS. The findings may be useful to further elucidate and drive the interpretation of the results highlighted in the main text of this manuscript, related to capabilities of AWS and TFA in predictive speech comprehension.

The behavioral facilitation in RTs found in previous studies (Gastaldon et al., 2020; Piai et al., 2014, 2015) is replicated in both groups: the HC condition induces faster responses, and the lexical frequency effect is present only in the LC condition, in which participants cannot retrieve the word before picture onset. Interestingly, the interaction between condition and group indicates that group differences are led by the HC condition: AWS are significantly slower than TFA controls, suggesting that they may be impaired in successfully initiating word production under constraining conditions.

Compatibly, when considering time-frequency analyses, the predictability of words to be produced elicited an evident alpha–beta power decrease associated to clusters in bilateral posterior frontal cortex, right sensorimotor cortices, and left inferior parietal cortex in AWS. On the other hand, in TFA, this effect was associated to a larger number of clusters, mainly in bilateral temporal and inferior parietal areas, right insula, right sensorimotor cortices, and right posterior frontal cortex.

When comparing AWS to TFA, the stuttering group displayed a stronger power decrease in the beta2 range in the left somatosensory cortex (in an earlier time-window, ~300 ms after the onset of the silent interval), in bilateral anterior frontal areas and in the right inferior frontal cortex (at a later time-window, ~500-700 ms after the onset of the silent interval). Additionally, a stronger decrease in the alpha range has been found in posterior frontal areas (pre-motor cortex and SMA complex), which emerged earlier in the left hemisphere (~0-200 ms after the onset of the silent interval) and later in the right hemisphere (~600-800 ms after the onset of the silent interval). Interestingly, an effect in the opposite direction is observed in the alpha-beta1 range in the right anterior frontal regions in the second half of the silent gap (~400-800 ms), whereby AWS display a reduced power decrease relative to TFA. Overall, this pattern may reflect an increased difficulty and an increased effort (thus triggering the recruitment of compensatory mechanisms) for AWS to prepare for production – possibly further modulated by altered executive control – in order to initiate and maintain fluent speech. Compatibly, the exaggerated power decrease in sensorimotor regions often reported in DS has been commonly interpreted as reflecting this increased difficulty and/or reduced automaticity/proficiency in initiating speech-motor programs (Mersov et al., 2016; Mock et al., 2016). This is also in line with suggestions advanced by computational simulations of stuttering, such as those based on the DIVA/GODIVA models (Chang & Guenther, 2020; Civier et al., 2010, 2013), showing that defective feedforward control in DS may easily result in motor-speech impairments, also biasing the system towards an overreliance on auditory feedbacks (Civier et al., 2010). In this context, the involvement of the right frontal cortex in stuttering has been commonly interpreted as underlying compensatory mechanisms (e.g. Neef et al., 2018, 2016; Preibisch et al., 2003). However, recent observations suggest that this exaggerated activity may also represent "maladaptive" compensations, or perhaps "pathological" activity, mainly related to altered executive and inhibitory mechanisms, thus speculatively resulting in stronger motor-speech difficulties for AWS (Neef et al., 2016, 2018). Interestingly, from observations during the debriefing after the experimental session, AWS participants did report increased difficulty in initiating the response when they could predict the word to produce, further suggesting that internally-generated and/or anticipated word production may result in additional burdening of their speech-motor system.

AWS show also differences in posterior areas. In the left temporal cortex, they showed a delayed and less marked power suppression in beta1 relative to controls, in an early time window of analysis. Given the region involved and the timing of the effect, we speculate that this may reflect a less efficient word selection process. In this context, a recent study suggested that a defective inhibitory control in lexical selection in AWS may contribute to DS, further destabilizing an already inefficient speech-motor system (Maxfield, 2020). Still in posterior regions, AWS also displayed a reduced power decrease in the beta range in the right inferior parietal cortex, possibly suggesting a weaker involvement of contextual model updating, in line with one of the roles proposed for this region in the context of an extended prediction network (Geng & Vossel, 2013; Siman-Tov et al., 2019).

When considering the correlation between stuttering and neural modulations, the presence of a negative correlation between beta1 power change in the left posterior frontal cortex (i.e., premotor cortex and SMA complex) and SSI-4 values supports the hypothesis that the increased engagement of the SMA complex may be related to disfluency severity, thus resulting in higher levels of “neural effort” and compensatory attempts (Mersov et al., 2016; Mock et al., 2016). As mentioned in the Discussion section of the main text, the SMA complex plays a role in the DIVA/GODIVA framework, specifically in the planning, sequencing, and implementation of the motor aspects of phonological representations, as well as in the coordinated and timed initiation of articulatory gestures. The present findings are compatible with the model, pointing towards the existence of an increased neural demand (and neural effort) on sequence planning in AWS during preparation for word production. In this context, the positive correlation highlighted between the facilitation in response times (more negative values corresponding to a greater facilitation) and beta2 power change in the right motor cortex of AWS corroborates the interpretation of increased activation in sensorimotor regions (reflected in stronger power decrease) as the result of the deployment of effective compensation strategies for initiating and maintaining fluent speech production (see Etchell et al., 2018, for a recent review of the literature in DS). Taken together, the present results about speech preparation for production in DS are in line with the previous evidence proposing the existence of “adaptive” or “maladaptive” compensatory mechanisms in AWS, mainly involving bilateral sensorimotor and frontal regions, in response to deficient speech-motor networks.

As an additional remark, albeit not central to the research questions of this study, we also note that the positive correlation between facilitation in response times and alpha power change in the left inferior parietal cortex of TFA supports the proposal put forward by Piai and colleagues (Piai et al., 2017, 2018), according to which left posterior areas are more prominently involved in the emergence of alpha–beta power decrease and of behavioral facilitation in context-induced word production.

**Supplementary references**

Busan, P. (2020). Developmental stuttering and the role of the supplementary motor cortex. *Journal of Fluency Disorders*, 105763. https://doi.org/10.1016/j.jfludis.2020.105763

Chang, S., & Guenther, F. H. (2020). Involvement of the Cortico-Basal Ganglia-Thalamocortical Loop in Developmental Stuttering. In *Frontiers in Psychology* (Vol. 10, p. 28). Frontiers Media S.A. https://doi.org/10.3389/fpsyg.2019.03088

Civier, O., Bullock, D., Max, L., & Guenther, F. H. (2013). Computational modeling of stuttering caused by impairments in a basal ganglia thalamo-cortical circuit involved in syllable selection and initiation. *Brain and Language*, *126*(3), 263–278. https://doi.org/10.1016/j.bandl.2013.05.016

Civier, O., Tasko, S. M., & Guenther, F. H. (2010). Overreliance on auditory feedback may lead to sound/syllable repetitions: Simulations of stuttering and fluency-inducing conditions with a neural model of speech production. *Journal of Fluency Disorders*, *35*(3), 246–279. https://doi.org/10.1016/j.jfludis.2010.05.002

Desikan, R. S., Ségonne, F., Fischl, B., Quinn, B. T., Dickerson, B. C., Blacker, D., Buckner, R. L., Dale, A. M., Maguire, R. P., Hyman, B. T., Albert, M. S., & Killiany, R. J. (2006). An automated labeling system for subdividing the human cerebral cortex on MRI scans into gyral based regions of interest. *NeuroImage*, *31*(3), 968–980. https://doi.org/10.1016/j.neuroimage.2006.01.021

Etchell, A. C., Civier, O., Ballard, K. J., & Sowman, P. F. (2018). A systematic literature review of neuroimaging research on developmental stuttering between 1995 and 2016. *Journal of Fluency Disorders*, *55*, 6–45. https://doi.org/10.1016/j.jfludis.2017.03.007

Gastaldon, S., Arcara, G., Navarrete, E., & Peressotti, F. (2020). Commonalities in alpha and beta neural desynchronizations during prediction in language comprehension and production. *Cortex*, *133*, 328–345. https://doi.org/10.1016/j.cortex.2020.09.026

Geng, J. J., & Vossel, S. (2013). Re-evaluating the role of TPJ in attentional control: Contextual updating? *Neuroscience and Biobehavioral Reviews*, *37*(10), 2608–2620. https://doi.org/10.1016/j.neubiorev.2013.08.010

Jenkinson, N., & Brown, P. (2011). New insights into the relationship between dopamine, beta oscillations and motor function. *Trends in Neurosciences*, *34*(12), 611–618. https://doi.org/10.1016/j.tins.2011.09.003

Jenson, D., Reilly, K. J., Harkrider, A. W., Thornton, D., & Saltuklaroglu, T. (2018). Trait related sensorimotor deficits in people who stutter: An EEG investigation of µ rhythm dynamics during spontaneous fluency. *NeuroImage: Clinical*, *19*, 690–702. https://doi.org/10.1016/J.NICL.2018.05.026

Joos, K., Ridder, D. De, Boey, R. A., & Vanneste, S. (2014). Functional connectivity changes in adults with developmental stuttering: A preliminary study using quantitative electro-encephalography. *Frontiers in Human Neuroscience*, *8*(OCT). https://doi.org/10.3389/fnhum.2014.00783

Kilavik, B. E., Zaepffel, M., Brovelli, A., MacKay, W. A., & Riehle, A. (2013). The ups and downs of beta oscillations in sensorimotor cortex. *Experimental Neurology*, *245*, 15–26. https://doi.org/10.1016/j.expneurol.2012.09.014

Lenth, R. (2020). *emmeans: Estimated Marginal Means, aka Least-Squares Means.* https://cran.r-project.org/package=emmeans

Maxfield, N. D. (2020). Inhibitory Control of Lexical Selection in Adults who Stutter. *Journal of Fluency Disorders*, 105780. https://doi.org/10.1016/j.jfludis.2020.105780

Mersov, A.-M., Jobst, C., Cheyne, D. O., & De Nil, L. (2016). Sensorimotor Oscillations Prior to Speech Onset Reflect Altered Motor Networks in Adults Who Stutter. *Frontiers in Human Neuroscience*, *10*, 443. https://doi.org/10.3389/fnhum.2016.00443

Mock, J. R., Foundas, A. L., & Golob, E. J. (2016). Cortical activity during cued picture naming predicts individual differences in stuttering frequency. *Clinical Neurophysiology*, *127*(9), 3093–3101. https://doi.org/10.1016/J.CLINPH.2016.06.005

Neef, N. E., Anwander, A., Bütfering, C., Schmidt-Samoa, C., Friederici, A. D., Paulus, W., & Sommer, M. (2018). Structural connectivity of right frontal hyperactive areas scales with stuttering severity. *Brain*, *141*(1), 191–204. https://doi.org/10.1093/brain/awx316

Neef, N. E., Bütfering, C., Anwander, A., Friederici, A. D., Paulus, W., & Sommer, M. (2016). Left posterior-dorsal area 44 couples with parietal areas to promote speech fluency, while right area 44 activity promotes the stopping of motor responses. *NeuroImage*, *142*, 628–644. https://doi.org/10.1016/J.NEUROIMAGE.2016.08.030

Piai, V., Klaus, J., & Rossetto, E. (2020). The lexical nature of alpha-beta oscillations in context-driven word production. *Journal of Neurolinguistics*, *55*, 100905. https://doi.org/10.1016/j.jneuroling.2020.100905

Piai, V., Meyer, L., Dronkers, N. F., & Knight, R. T. (2017). Neuroplasticity of language in left-hemisphere stroke: Evidence linking subsecond electrophysiology and structural connections. *Human Brain Mapping*, *38*(6), 3151–3162. https://doi.org/10.1002/hbm.23581

Piai, V., Roelofs, A., & Maris, E. (2014). Oscillatory brain responses in spoken word production reflect lexical frequency and sentential constraint. *Neuropsychologia*, *53*(1), 146–156. https://doi.org/10.1016/j.neuropsychologia.2013.11.014

Piai, V., Roelofs, A., Rommers, J., & Maris, E. (2015). Beta oscillations reflect memory and motor aspects of spoken word production. *Human Brain Mapping*, *36*(7), 2767–2780. https://doi.org/10.1002/hbm.22806

Piai, V., Rommers, J., & Knight, R. T. (2018). Lesion evidence for a critical role of left posterior but not frontal areas in alpha–beta power decreases during context-driven word production. *European Journal of Neuroscience*, *48*(7), 2622–2629. https://doi.org/10.1111/ejn.13695

Preibisch, C., Neumann, K., Raab, P., Euler, H. A., Von Gudenberg, A. W., Lanfermann, H., & Giraud, A. L. (2003). Evidence for compensation for stuttering by the right frontal operculum. *NeuroImage*, *20*(2), 1356–1364. https://doi.org/10.1016/S1053-8119(03)00376-8

R Core Team. (2018). R: A language and environment for statistical computing. R Foundation for Statistical Computing. http://www.r-project.org

Roux, F., Armstrong, B. C., & Carreiras, M. (2017). Chronset: An automated tool for detecting speech onset. *Behavior Research Methods*, *49*(5), 1864–1881. https://doi.org/10.3758/s13428-016-0830-1

Saltuklaroglu, T., Bowers, A., Harkrider, A. W., Casenhiser, D., Reilly, K. J., Jenson, D. E., & Thornton, D. (2018). EEG mu rhythms: Rich sources of sensorimotor information in speech processing. *Brain and Language*, *187*, 41–61. https://doi.org/10.1016/J.BANDL.2018.09.005

Siman-Tov, T., Granot, R. Y., Shany, O., Singer, N., Hendler, T., & Gordon, C. R. (2019). Is there a prediction network? Meta-analytic evidence for a cortical-subcortical network likely subserving prediction. *Neuroscience & Biobehavioral Reviews*, *105*, 262–275. https://doi.org/10.1016/j.neubiorev.2019.08.012
